# Single-cycle SARS-CoV-2 vaccine elicits high protection and sterilizing immunity in hamsters

**DOI:** 10.1101/2023.05.17.541127

**Authors:** Martin Joseph Lett, Fabian Otte, David Hauser, Jacob Schön, Enja Tatjana Kipfer, Donata Hoffmann, Nico J. Halwe, Lorenz Ulrich, Yuepeng Zhang, Vladimir Cmiljanovic, Claudia Wylezich, Lorena Urda, Christopher Lang, Martin Beer, Christian Mittelholzer, Thomas Klimkait

## Abstract

Vaccines have been central in ending the COVID-19 pandemic, but newly emerging SARS-CoV-2 variants increasingly escape first-generation vaccine protection. To fill this gap, live particle-based vaccines mimicking natural infection aim at protecting against a broader spectrum of virus variants. We designed “single-cycle SARS-CoV-2 viruses” (SCVs) that lack essential viral genes, possess superior immune-modulatory features and provide an excellent safety profile in the Syrian hamster model. Full protection of all intranasally vaccinated animals was achieved against an autologous challenge with SARS-CoV-2 virus using an Envelope-gene-deleted vaccine candidate. By deleting key immune-downregulating genes, sterilizing immunity was achieved with an advanced candidate without virus spread to contact animals. Hence, SCVs have the potential to induce a broad and durable protection against COVID-19 superior to a natural infection.

## Introduction

Since its first appearance in 2019, SARS-CoV-2 has spread rapidly worldwide and continues to circulate in many countries, causing symptoms and COVID-19 disease, despite an unprecedented, quick deployment of effective first-generation mRNA- and vector-based vaccines (Baden et al., 2021; Beesley et al., 2023; Polack et al., 2020; Sadoff et al., 2022; Voysey et al., 2021), targeting the viral Spike (S) protein. Since then, multiple virus variants have emerged, carrying escape mutations mainly in the S gene that correlate with declining protection rates (Muik et al., 2022; Tseng et al., 2022).

To combat new variants of the virus and induce an immune response to additional viral proteins, recent vaccine approaches focus on attenuating the virus (Nouailles et al., 2023; Wang et al., 2021) and on intranasal applications for stronger induction of mucosal immunity (Afkhami et al., 2022). One principal drawback of attenuated viral vaccines is the residual risk of an accidental reversion to virulence, i.e., causing the wild-type like disease from which one would like to protect (Minor, 1992; Platt et al., 2014). This aspect is particularly crucial for key risk groups: immunocompromised, transplanted and elderly people, or cancer patients.

To generate a safe but effective SARS-CoV-2 vaccine with improved properties inducing a similarly broad immune response as live SARS-CoV-2 viruses, we designed a ‘single-cycle infection concept’. The deletion of one essential structural gene from the viral genome, combined with a stable cellular trans-complementation system as used for other coronaviruses (Almazan et al., 2013; Gutierrez-Alvarez et al., 2021; Netland et al., 2010; X. Zhang et al., 2021), leads to the production of intact but propagation-defective particles that may serve as SARS-CoV-2 vaccine candidates. We opted to eliminate the poorly immunogenic Envelope (E) gene and inserted an eGFP reporter in the reading frame of E (ΔE^G^).

In addition, we deleted two of the accessory genes described to be crucial for down-modulating the anti-viral defense (Kimura et al., 2021; Yoo et al., 2021; Y. Zhang et al., 2021), creating a triple-deletion virus termed ΔE^G^68 (Fig. 1A).

Eliminating these SARS-CoV-2 accessory proteins, encoded e.g. by open reading frames (ORFs) ORF3a, ORF6, ORF7a and ORF8 (Silvas et al., 2021), is expected to increase the immunogenicity of single-cycle viruses beyond that of a natural SARS-CoV-2 infection while retaining their safety, which is mainly based on the E gene deletion. In addition, ORF6 has been described to suppress T cell responses (Yoo et al., 2021) and eliminate the interferon (IFN) response in the infected cell (Kimura et al., 2021). ORF8 had been shown to reduce the T-cell response *in vivo* (Y. Zhang et al., 2021).

This study thoroughly investigates the properties of a single-cycle, triple-deletion vaccine virus (ΔE^G^68) and assesses the direct impact of eliminating ORF6 and ORF8 by comparing it to an “E-deleted only” vaccine virus (ΔE^G^). We show evidence for enhanced immune stimulation, the elicitation of full protection against challenge infection, and for sterilizing immunity in the Syrian hamster model.

## Results

### Single-cycle virus stability and in vitro safety profile

Both SCV candidates were obtained using the ISA-based method described previously (Fig. 1A, fig. S1A, from design to vaccine virus in ∼4 weeks) (Kipfer et al., 2023; Melade et al., 2022). ΔE^G^68 and ΔE^G^ were efficiently rescued in E-complementing HEK293T cells (HEK293T-indE) and propagated in a Vero E6-based cell line stably expressing the E protein (VeroE2T). The presence and functionality of E in the established cell line were assessed by mRNA detection (fig. S1, B and C) or by virus complementation and propagation of cell-free progeny virus (Fig. 1B, fig. S1D and fig. S2). SCVs were monitored by antigen quick-tests (Kipfer et al., 2023) and quantified in focus formation assays (FFA).

**Figure 1:**
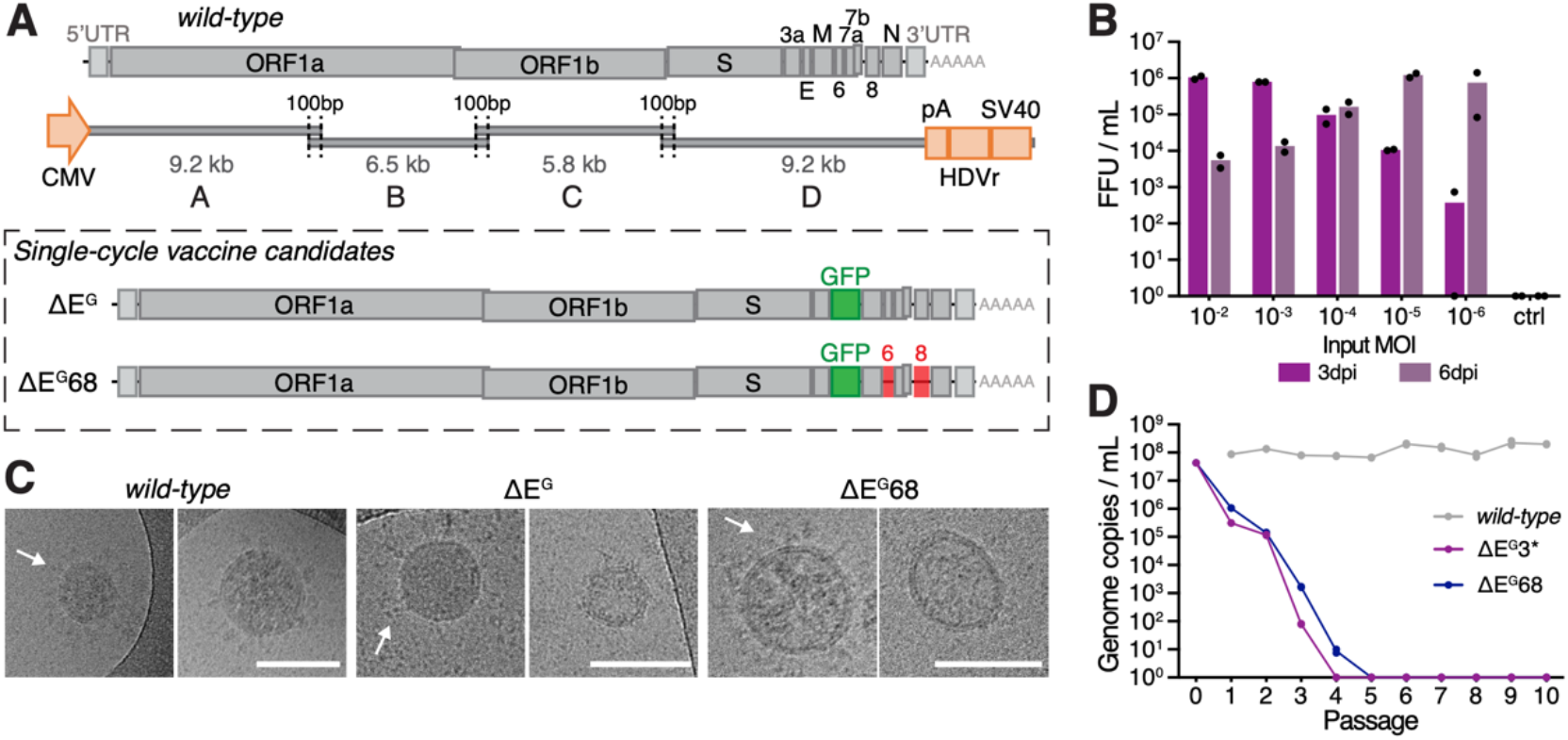
Single-cycle vaccine concept and viral characterization. **(A)** Schematic illustrating the SARS-COV-2 genomic landscape and the deletions/substitutions in ΔE^G^/ΔE^G^68, main structural and accessory proteins indicated. Four overlapping fragments covering the whole SARS-CoV-2 genome were amplified by PCR (Fragments A-D). The heterologous CMV promoter was cloned upstream of the 5’ UTR and a poly(A) tail, HDV ribozyme, and SV40 termination signal downstream of the 3’ UTR. **(B)** Complementation efficiency of VeroE2T cells, analyzed by FFA (focus forming assay) of ΔE^G^ infection at different MOI or medium-only control (ctrl) after 3 and 6 dpi (n=2 individual cultures), for corresponding genome copies see fig. S1D. **(C)** Transmission electron microscopy analysis of recombinant wild-type SARS-CoV-2 (rCoV2) or vaccine candidates ΔE^G^ and ΔE^G^68 showing the presence of the characteristic spike protein (indicated with arrows). **(D)** Passaging of 1:10 and 1:100 (after p2) dilutions of cell-free supernatant (Input = Passage 0) of wild-type SARS-CoV-2 (Muc-1, B.1), ΔE^G^ and ΔE^G^68 on non-complementing VeroE6 cells starting with an initial infection with an MOI of 1. Data from one representative experiment are shown; analysis was performed in duplicates. Scale bar is 100nm in (**C**)

The precise deletion of the three intended genes in vaccine virus candidate ΔE^G^68 and of the E-gene in ΔE^G^ and their stable functional elimination were verified after repeated passage in VeroE2T cells by NGS and Sanger sequencing . For candidate ΔE^G^, a several log-fold increase of viral loads was observed upon repeated passaging, attributable to a spontaneous frame shift mutation in ORF3a that introduced a translational stop codon. Thus, for high multiplicity of infection (MOI) experiments and *in vitro* safety passaging, ΔE^G^3* (ΔE^G^ with additional translational stop codon in ORF3a) was used. For animal safety data, the ΔE^G^ candidate was tested.

In order to demonstrate that SCVs indeed represent authentic viral particles that package the defective genome, virions were analyzed by transmission electron microscopy, which confirmed the efficient production of spike-carrying spheres with the expected size of 80-100 nm typical for SARS-CoV-2 virions. To assess lower levels of viral S protein observed on the vaccine candidates (Fig. 1C), surface labeling of cells infected with SCVs or *wild-type* control was performed. Vaccine candidates show a strong S-signal at cell-to-cell interfaces compared to a more clustered staining of cells infected with *wild-type*. This indicates differences in viral assembly and particle formation (fig. S1E) (Cattin-Ortola et al., 2021).

The single-cycle nature of the genetically modified vaccine candidates ΔE^G^68 and ΔE^G^ was demonstrated by infecting standard VeroE6 cells that are commonly used for SARS-CoV-2 propagation: Even after a high MOI infection, detectable virus of either candidate quickly vanished from the culture supernatant, in contrast to *wild-type* infections during passaging (Fig. 1D). The possible emergence of viral revertants at sub-detection levels in VeroE6 cells was excluded by inoculating the producer cell line VeroE2T for 6 days with supernatant samples from passages 1 to 10. As 1-5 focus-forming units (FFUs) are sufficient to initiate full viral amplification on VeroE2T cells (Fig. 1B), an efficient propagation even of low-level revertants or newly emerging replicative viral variants would have been detected. None of the passages below 100 genomic copies/mL led to any rescuable replicative virus (fig. S2).

### Molecular characterization of vaccine candidates *in vitro*

We analyzed viral protein expression in infected VeroE6-TMPRSS2 cells (stable expression of TMPRSS2 in VeroE6 cells, fig. S1B) by immunoblotting and immunocytochemistry. As additional control we included the E-defective mutant E**^fs^ (two back-to-back stop codons (*) and insertion of an additional G-nucleotide (frameshift (fs)) after the first 7 amino acids of E) that retains the RNA sequence and secondary structure to a large extent. At 24h post-infection, similar viral protein levels were found as for *wild-type* virus (Fig. 2, A and B). The expression of NSP2 (non-structural protein 2, as reference for virus input), Nucleocapsid protein (N) and S was comparable, with elevated levels of cleaved S (subunit S2) only for the E**^fs^ mutant. For ΔE^G^68, absence of ORF6 and ORF8 was confirmed, while ORF7 as interjacent gene remained expressed in all tested variants. Immunoblot data also confirmed the expected ORF3a truncation in ΔE^G^3* due to the translational stop codon (Fig. 2A).

In summary, we observed for both ΔE^G^68 and ΔE^G^ vaccine candidates a close-to-*wild-type* expression level of all structural components, similar particle properties, and strict single-cycle infection in standard cells. This molecular characterization led us to verify the immunizing performance of our SC-vaccines *in vitro* and *in vivo*.

**Figure 2:**
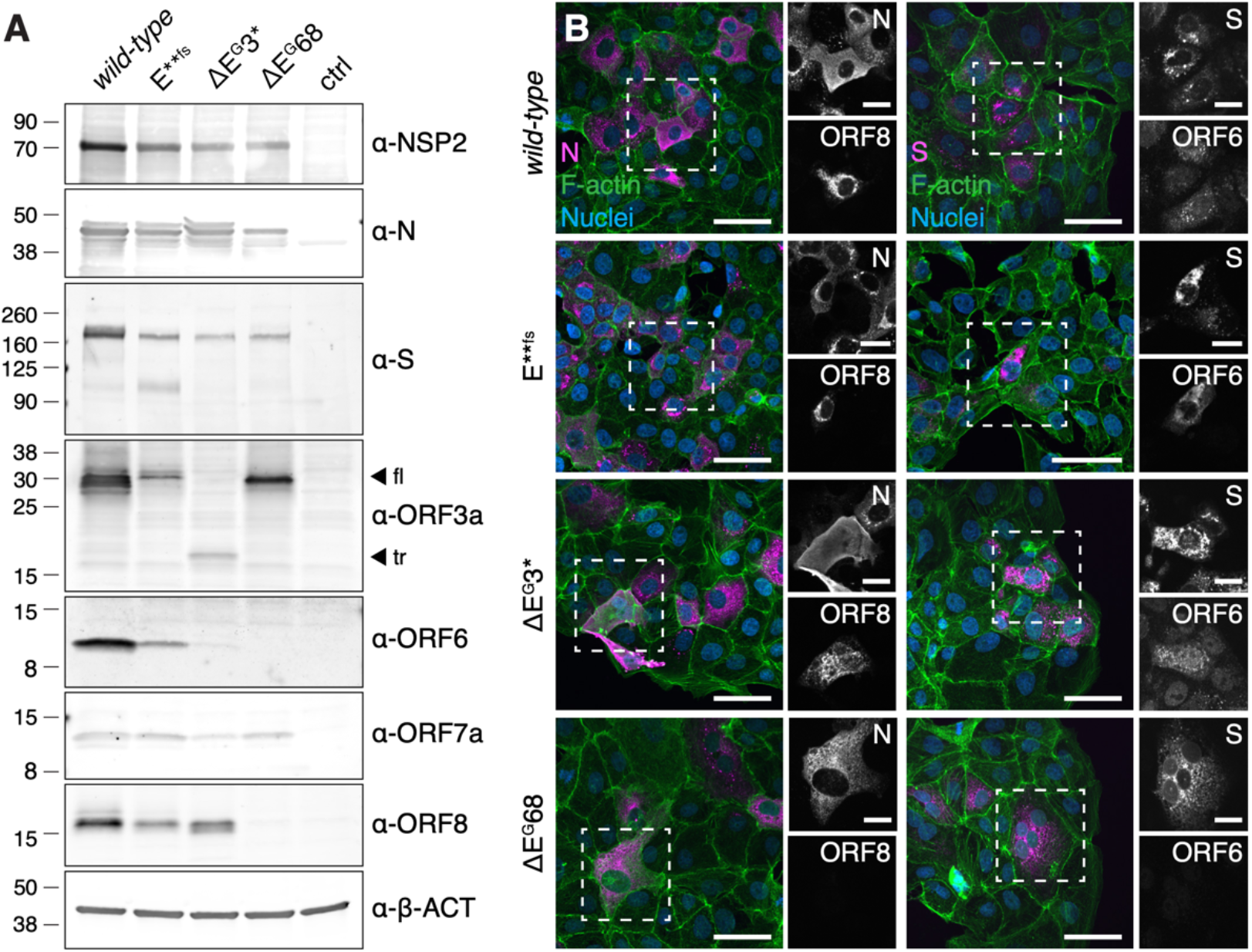
Evaluation of protein synthesis after infection with mutant virus. **(A)** Immunoblot analysis of viral protein production in VeroE6-TMPRSS2 cells infected with rCoV2, E**fs, ΔE^G^, ΔE^G^68 or medium only (ctrl), probed with anti-NSP2, anti-N, anti-S, anti-ORF3a (full-length (fl) and truncated (tr) forms indicated with arrows), anti-ORF6, anti-ORF7a, anti-ORF8 and anti-beta-actin (β-ACT) antibodies. **(B)** Detection of N and S (magenta), F-actin (green), nuclei (blue) and ORF6 or ORF8 in VeroE6-TMPRSS2 cells infected with rCoV2, E**fs, ΔE^G^ or ΔE^G^68. Scale bar is 50µm and 20µm (overview and ROI images, respectively)

### *In vitro* immuno-modulatory responses to vaccine candidates

Immune-downmodulating functions have been reported for ORF6 and ORF8 and to a lesser extent for the Envelope protein (Chen et al., 2022; Kimura et al., 2021; Vann et al., 2022; Yoo et al., 2021; Y. Zhang et al., 2021). To test whether the lack of E, ORF6 and ORF8 in the SCV could provide a stronger immune response than *wild-type* virus, we transiently expressed each gene in monocytic THP-1 cells as a model for antigen-presenting cells (APC). The impact of the newly introduced protein on immunological markers was then assessed by cell surface staining of antigen-presenting proteins (HLA-A/B/C, HLA-DR), the co-stimulators CD40, CD44, CD70, CD80 and CD275, and complement cascade protein (CD59). At 48 hours post-transfection, we observed a downregulation of CD80 and CD275 on THP-1 cells for all three proteins compared to a control plasmid (Fig. 3, A to C), while no change was observed in the expression of HLA-DR and CD70, thus excluding labeling artefacts (Fig. S3). The direct effect on the HLA seems more modest (Fig S3B). Taken together, these data indicate that the expression of ORF6, ORF8 and E correlates with a diminished presentation capacity on APCs. We then infected alveolar basal epithelial cells (A549) for 24 hours and stained them with the same panel, excluding HLA-DR. Two different SARS-CoV-2 strains served as controls: the original Wuhan strain (B.1), which is the basis of our mutants, and the recent Omicron XBB.1.5 strain, which naturally contains a premature stop codon at position 8 of ORF8, i.e., loss of ORF8 function as a result of natural selection (fig. S4). A549 cells downregulated HLA-A/B/C and CD275 when infected with the Wuhan strain, but not with ΔE^G^68, whereas Omicron XBB.1.5 and E**^fs^ show only partial down-regulation, evoking a role of ORF8 (Fig 3, D and E).

Culture supernatants from infection of A549 or HEK293T cells were incubated with non-infectable THP-1 cells for 48 hours before staining (fig. S3). Of interest, for HLA-A/B/C and CD80 we observed the same effect of the deletion as seen in overexpression experiments: while receptor expression was downregulated by *wild-type* infection, ΔE^G^68 SCV conversely induced a higher expression (Fig. 3, F to H). The E**^fs^ mutant displayed intermediate expression levels, suggesting additive, non-overlapping functions of ORF6, ORF8 and E. The observation that the effect was seen on both infected and non-infectable cells suggests that these ORFs directly and indirectly impaired antigen presentation.

**Figure 3:**
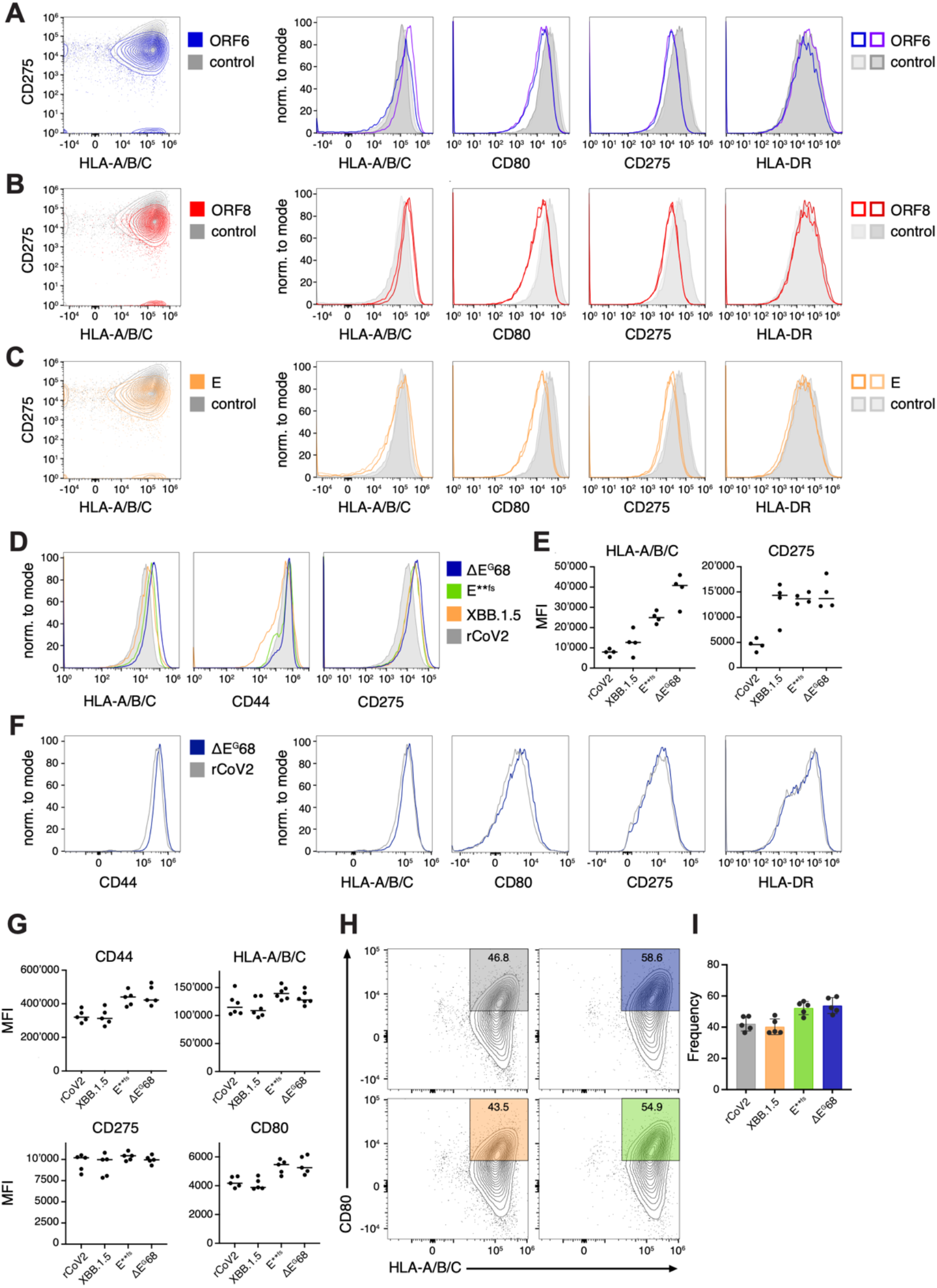
Immunomodulation by E, ORF6 and ORF8 proteins. (**A-C**) Modulation after transfection: Flow cytometry staining of THP-1 cells for HLA-A/B/C, CD80, CD275, and HLA-DR surface expression 48h after transfection with expression plasmids for ORF6 (**A**), ORF8 (**B**), or Envelope (**C**) proteins, compared with control transfection. (**D-I**) Modulation after infection: A549-ACE2-TMPRSS2 cells were infected with rCoV2, E**fs, ΔE^G^68, or XBB.1.5 SARS-CoV-2 virus (MOI = 0.1) for 24h, and stained for HLA-A/B/C, CD44 and CD275. Median fluorescence intensity (MFI) of HLA-A/B/C and CD274 is shown (**E**). The same infection was conducted on HEK293T and their respective supernatant is then applied on THP-1 for 48h before surface staining and analysis. (**F**) Histogram showing the expression of CD44, HLA-A/B/C, CD80, CD275, and HLA-DR on THP-1 after 48 h. (**G**) Median fluorescence intensity of HLA-A/B/C, CD80, and CD275 marker on THP-1 after 48 h incubation. The downregulation of the HLA and co-molecule can be seen when full-length virus is used as seen in the dot plot (**H**) comparing wild-type or ΔE^G^68 condition for their expression of CD80 and HLA-A/B/C. The frequency of cells outside of the gate in (**H**) is shown in (**I**).

### Vaccination and challenge infection in the Syrian hamster model

Our single-cycle vaccine concept was examined *in vivo* using the highest achievable dose for ΔE^G^68 or a low dose for the construct ΔE^G^. Candidates or controls were administered intranasally to 5-to 6-week-old Syrian hamsters, an infection model used for safety and efficacy due to efficient viral spread (Sia et al., 2020). Naïve contact hamsters were co-housed with immunized hamsters 24 hours after vaccine application, to be separated again for 24 hours only immediately before boost immunization or challenge infection (Fig. 4A).

Hamsters were immunized with 2.4*10^4^ FFUs of ΔE^G^68 (n=12) or 3.5*10^2^ FFUs of ΔE^G^ (n=8) in 100 µL per animal (fig. S5). Following immunization, all animals continually gained weight as expected (Fig. 4, B and C). A minimal ’dip’ in mean body weight on days 2-3 was observed in all experimental groups, including contact animals (Fig. 4B), and is typical for and attributable to procedural stress. Since body weight loss usually occurs when Syrian hamsters are inoculated with *wild-type* SARS-CoV-2, as shown by subsequent challenge infection of sham-treated animals (Fig. 4D), this indicates that both vaccine candidates were very well tolerated.

Already 19 days post immunization (dpim), profound SARS-CoV-2 specific humoral immune responses were confirmed in all animals vaccinated with ΔE^G^68 or ΔE^G^ and were even more pronounced after the second immunization (33 dpim) (Fig. 4E).

The singe-cycle nature of ΔE^G^68 and ΔE^G^ was confirmed by rapidly declining viral RNA signals at 3 dpim (10^4,7^ or 10^3.8^ mean genome copies/mL, resp.) and 7 dpim (10^2^ or 10^2.1^ genome copies/mL, resp.) after prime-immunization. This was close to or below the applied threshold of the assay used (Fig. 4F, grey area).

On 3 dpim, two ΔE^G^68 contacts became positive with a mean of 10^3.5^ genome copies/mL (Fig. 4F, light blue) as compared to the input of 2*10^6^ RNA copies administered per animal. An E-gene-specific RT-qPCR assay (Corman et al., 2020) verified the deletion of E and excluded the possibility of a reversion, which was further strengthened by the lack of clinical signs and the absence of any further virus spread (table S2 or Fig. 4, B to D and F). The low virus RT-qPCR signal observed at 7 dpim for a single ΔE^G^68 contact hamster was correlated with the presence of SARS-CoV-2 specific antibodies on day 19 (Fig. 4E, light blue). However, based on the way of sampling, we cannot exclude that during nasal washing, some tissue cells might have been aspirated causing increased variance on 3 and 7 dpim.

**Figure 4.**
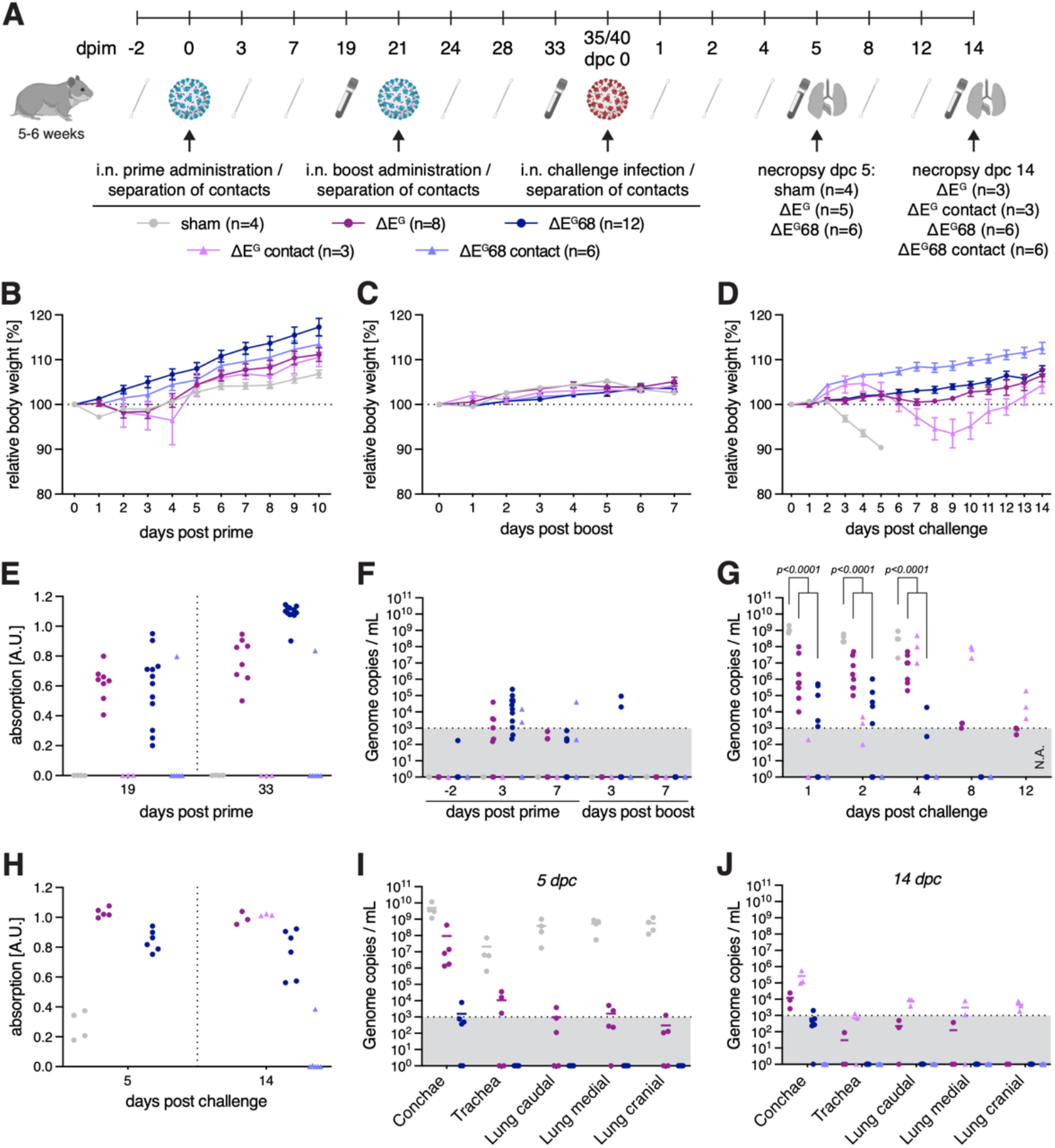
Immunization of Syrian hamsters with ΔE^G^ and ΔE^G^68 live vaccine candidates and challenge infection. **(A)** Experimental setup and timeline including a prime-boost-immunization and subsequent virus challenge. At indicated time points serum and nasal washing samples were taken. Organ samples were obtained on the days of necropsy. Serum samples were used to detect SARS-CoV-2 RBD (receptor binding domain)-specific antibodies by ELISA or neutralizing antibodies. Genomic RNA loads in nasal washings and organ samples were investigated by SARS-CoV-2 polymerase gene-specific RT-qPCR. (**B-D**) Relative body weight after intranasal prime (**B**), boost immunization (**C**) and challenge infection (**D**). (**E**) Humoral immune response after prime and boost immunization (dpim 19 and 33, resp.), determined by ELISA against the SARS-CoV-2 RBD of S. (**F, G**) Virus genome copy numbers detected in nasal washing samples following prime and boost immunization (**F**) and challenge infection (**G**) (note: no data available for ΔE^G^68 and ΔE^G^68 contact animals at 12 dpc in (**G**)). (**H**) Humoral immune response after challenge (5 and 14 dpc), determined by SARS-CoV-2 RBD specific ELISA. (**I, J**) Viral genome copies in organ samples 5 dpc (**I**) and 14 dpc (**J**). Mean and S.E.M. (**B, C** and **D**), scatter plots (**I** and **J**) show mean values as line, two-way anova followed by Bonferroni’s test (**G**).

After boost immunization, two out of 12 animals immunized with ΔE^G^68 had an RNA signal on day 3. On day 7, no vaccine RNA remained detectable in any of the ΔE^G^ or ΔE^G^68 immunized or contact animals (Fig. 4F).

Following homologous challenge with *wild-type* SARS-CoV-2 virus (∼10^2.5^ TCID, Wuhan B.1), no weight loss was observed in the ΔE^G^68-and ΔE^G^-vaccinated groups, while all sham-vaccinated animals lost body weight until 5 days post challenge infection (dpc) (Fig. 4D). Moreover, only very low viral loads, close to the threshold of quantification (grey area in Fig. 4G), were recovered from nasal washes on days 1, 2 and 4 after challenge infection of the ΔE^G^68 vaccinated animals. This was in sharp contrast to and significantly different from the situation in sham-vaccinated animals (p<0.0001 for all three time points, Fig. 4G), in which 10^7^-10^9^ copies/mL were recovered. No viral genomes in nasal washing samples and no weight loss were observed in any of the 6 contact animals post challenge infection (Fig. 4G). This complete protection of the 6/6 contact animals strongly supports the notion of a sterilizing immunity achieved by the ΔE^G^68 SC-vaccine.

It is interesting to note that the high levels of a pre-challenge antibody response did not further increase following challenge infection. This argues for a full response with maximal antibody induction already during the boost immunization phase, leading to a strong mucosal replication block of the challenge virus (Fig. 4H).

Upon detailed organ examination of ΔE^G^68 immunized animals 5 dpc, a low viral load near the quantification limit was restricted to the nasal respiratory tract (grey area in Fig. 4I). On day 14 post challenge, the RNA levels in the conchae of ΔE^G^68-vaccinated animals were undetectable or below a quantifiable level. No signal was detected in the trachea or lungs of any of the animals (Fig. 4J). This indicates a high level of protection achieved with the SC vaccines.

For ΔE^G^, RT-qPCR revealed quantifiable viral loads only in the conchae, calculated to be at least 50-fold lower than in the sham-immunized animals and a nearly complete protection from virus replication was confirmed in lung tissues (Fig. 4I). On day 14 dpc, the RT-qPCR signal in the conchae and the lower respiratory tract was greatly reduced. Furthermore, none of the animals had experienced weight loss during this period (Fig. 4D).

When evaluating the weight of the contact animals of both vaccines, weight loss in the ΔE^G^ contact animals was greatly delayed compared to infected controls (Fig. 4D). There was no indication until 3 dpc, before a slight weight reduction of 6.5% occurred until 9 dpc. Infection controls had lost on average 9.6% in weight by day 5, when they were sacrificed for sampling. Weight loss in contact animals can be explained by reduced virus shedding after challenge for ΔE^G^ immunized animals, which was significantly lower than in sham-immunized controls (p<0.0001 for dpc 1, 2 and 4, Fig. 4G). Viral shedding by ΔE^G^ contact animals began at 2 dpc and was still detectable 12 dpc (Fig. 4G). At 4 dpc, the onset of prominent virus replication in the contact hamsters by far exceeded the shedding levels of the vaccinated animals.

Neutralizing antibody responses were quantified against Wuhan (B.1). In 10 out of 12 ΔE^G^68 vaccinated hamsters, neutralizing antibodies were already detectable after boost immunization (mean 1:229 for 100% neutralization dose) and remained stable after challenge for all vaccinated animals (5 dpc, 1:220; 14 dpc, 1:140) (table S3). For the ELISA-positive contact animal, a weak antibody response was detected on 33 dpim (1:40). For animals vaccinated with ΔE^G^, neutralizing antibodies were detected after challenge infection (5 dpc, 1:404; 14 dpc, 1:295) (table S4). Notably, it can’t be excluded that neutralization for ΔE^G^ would score positively before challenge, as the obtained serum volume was technically limiting to assess lower dilutions. Only one in four sham animals had a low titer (1:20), ruling out that the rise to neutralizing antibodies was based on the virus challenge. For all ΔE^G^ contact animals, a very low neutralization titer was apparent at 14 dpc (table S4).

Taken together, both vaccine candidates elicit high protection in the Syrian hamster. In addition, the deletion of ORF6 and ORF8 led to a sterilizing immunity in all vaccinated animals potentially due to a stronger immune response by IFN-mediated signaling, improved immune stimulation and/or higher vaccine inoculum.

## Discussion

Efficient vaccines must have key properties to generate an immune response. First, providing or generating enough targets recognized by host antibodies, and second, inducing sufficient activation of T lymphocytes. In addition to strong immunogenicity, it is essential to ensure maximum safety. The two vaccine candidates reported here combine these properties. Our single-cycle vaccine generates *wild-type* like viral particles, which induce an accumulation of viral proteins in the host cell, serving as targets for B and T cells. This implies that efficient replication of viral RNA has occurred. Deletion of ORF6 and ORF8, two anti-inflammatory proteins that antagonize T cell activation, further supports a strong host response as suggested by our *in vitro* data and published literature (Cattin-Ortola et al., 2021; Chen et al., 2022; Kimura et al., 2021; Yoo et al., 2021; Y. Zhang et al., 2021).

We show that our candidate ΔE^G^68 causes higher surface expression of HLA molecules and co-stimulatory factors on infected cells or surrounding APCs, in particular CD80 (B7-1) and CD275 (B7-H1/ICOSLG), both involved in T cell stimulation (Wong et al., 2003; Yu et al., 2000). Notably, humans with a defective CD275 gene produce low levels of IgG, IgA, and memory B cells (Roussel & Vinh, 2021). The measurable effect on infected cells, but also on non-infectable cells in contact, suggests an indirect effect through local inflammation. These elements argue for greater immunogenicity of the SCV compared to its native counterpart.

Maximum safety of our vaccine approach is ensured by the demonstrated single-cycle concept. This prevents viral propagation, and unlike an attenuated virus approach, which relies on the immune system to combat a weakened virus, could enable the use in immunocompromised people.

Furthermore, we achieved sterilizing immunity for ΔE^G^68 in Syrian hamsters, a characteristic that is fundamental to preventing viral spread in humans and that has not been achieved in other vaccine candidates so far (Jung et al., 2022; Martinez-Baz et al., 2023). This might be due to enhanced local immunity after nasal application, which prevents viral shedding (Miteva et al., 2022). Interestingly, we observed transmission of high dose ΔE^G^68 vaccine to one of six contact animals. Spontaneous genetic reversion was excluded by RT-qPCR. No further propagation or weight loss was observed, indicating a passive transfer of the vaccine virus. The transfer was accompanied by seroconversion, implying that even a very small dose of SCV is sufficient to induce a high serological response.

It should be mentioned here that we had to repeat one SARS-CoV-2 challenge infection due to an erroneous over-dilution with no detection of infectious challenge virus (see Materials and Methods). However, all experimental data confirm that this had no influence on the overall results and that the repeated challenge infection could be classified as valid.

Taken together, our proposed single-cycle vaccine concept consolidates the high safety of an intranasally applied vaccine that induces sterilizing immunity, which will be key to overcoming the ongoing SARS-CoV-2 outbreaks.

## Materials and methods

### Animal experiments

#### Model

All procedures involving animals were evaluated by the responsible ethics committee of the State Office of Agriculture, Food Safety, and Fishery in Mecklenburg– Western Pomerania (LALLF M-V) and gained governmental approval under the registration numbers LVL MV TSD/ 7221.3-1-041/20. Specific pathogen free male Syrian hamsters (*Mesocricetus auratus*) (Janvier labs, RjHan:AURA) were kept at 20 to 22°C and a relative humidity of 45 ± 10% on a 12-hour light/dark cycle, fed with commercial rodent chow (Ssniff, Soest, Germany), and provided with water ad libitum. Age of the animals at prime immunization is 5 weeks for ΔE^G^ and 6 weeks for ΔE^G^68. Generally, hamsters underwent a daily physical examination and bodyweight routine.

#### ΔE^G^ immunization

Eight hamsters were intranasally inoculated with 100 µL of ΔE^G^ virus stock (3.5*10^3^ FFU, fig. S5) at day 0 and boosted with the same dose at day 21. Four hamsters were inoculated with 100 µL of supernatant from uninfected cells and therefore served as sham vaccinated controls. The three direct contact animals were co-housed with ΔE^G^ immunized animals, but were separated for 24 hours just prior to immunizations and challenge, respectively. Nasal washing samples were taken at day -2, 3, 7, 24, 28, 36, 37, 39, 43 and 47 days post immunization (dpim), by applying 200 µL of PBS into each nostril and collecting the reflux under short isoflurane inhalation anesthesia. Serum samples were taken by puncturing the *V. saphena* at 19 and 33 dpim for serological evaluation. At 35 dpim eight ΔE^G^ immunized animals and four sham vaccinated control animals were challenged by intranasal inoculation using 10^2,5^ TCID_50_/animal of SARS-CoV-2 virus (Wuhan-1, GenBank No. MT108784 (*34*)) in a 70 µL volume (calculated from back-titration). Five days post challenge (dpc), five ΔE^G^ immunized hamsters and the sham vaccinated control hamsters were sacrificed and sera or organ samples from upper and lower respiratory tract were collected during necropsy. 14 dpc three ΔE^G^ immunized hamsters and three contact animals were euthanized and serum sample as well as organ samples from upper and lower respiratory tract were collected during necropsy.

#### ΔE^G^68 immunization

Twelve hamsters were intranasal inoculated with 100 µL of ΔE^G^68 virus stock (2.4*10^5^ FFU, fig. S5) at day 0 and boosted with the same dose at day 21. Six direct contact animals were co-housed with ΔE^G^68 immunized animals, but were separated for 24 hours prior to immunizations and challenge infection, respectively. Nasal washing samples were taken at dpim -2, 3, 7, 24, 28, 36, 37, 38, 41 (dpc1), 42 (dpc2), 44 (dpc4) and 48 (dpc8) by applying 200µL of PBS in each nostril and collecting the reflux under short isoflurane inhalation anesthesia. Serum samples were taken by puncturing the *V. saphena* at 19 and 33 dpim for serological evaluation. At 35 dpim the ΔE^G^68 immunized animals were inoculated using a miscalculated low dosage of SARS-CoV-2 virus (Wuhan-1, GenBank No. MT108784 (*34*)) with less than 1 TCID_50_/animal. The viral genome copies in this misdiluted inoculum were determined by RT-qPCR (RNA-dependent RNA polymerase (IP4) as target (*35*)) with a Ct-value of 35.64, representing 1089 genome copies/mL. With this highly diluted inoculum, we were unable to perform an endpoint titration and to initiate a productive infection when 70 µL of pure inoculum were applied to VeroE6 cells (0.32 cm^2^, n=7). Additionally, nasal washing samples were taken from all animals on the first three days after inoculation and were all negative by RT-qPCR (table S5). Therefore, a second challenge infection was performed with the same animals at 41 dpim applying 70 µL with 10^2.3^ TCID_50_/animal (Wuhan-1, GenBank No. MT108784 (*34*)), calculated from back-titration. Five days post challenge infection, six ΔE^G^68 immunized hamsters were euthanized and serum samples as well as organ samples from the upper and lower respiratory tract were collected during necropsy. 14 dpc six ΔE^G^ immunized hamsters and their respective six matching contact animals were euthanized and serum sample as well as organ samples from upper and lower respiratory tract were collected during necropsy.

#### Analysis of hamster samples

RNA from nasal washings and organ samples was extracted using the NucleoMag® VET Kit (Macherey-Nagel, Düren, Germany) in combination with a Biosprint 96 platform (Qiagen, Hilden, Germany). Viral RNA genomes were detected and quantified by real-time RT-qPCR on a BioRad real-time CFX96 detection system (BioRad, Hercules, USA). The target sequence for amplification was viral RNA-dependent RNA polymerase (IP4) (*27, 35*). Genome copies per mL sample were calculated based on a quantified standard RNA, where absolute quantification was done by the QX200 Droplet Digital PCR System in combination with the 1-Step RT-ddPCR Advanced Kit for Probes (BioRad, Hercules, USA). The detection limit was calculated to be 1000 copies per reaction. Serum samples were analysed using an indirect multispecies ELISA against SARS-CoV-2 RBD (*36*). Briefly, RBD coated plates or those treated with coating buffer-only were blocked with 5% skim milk in phosphate-buffered saline, pH 7.5. Serum samples were incubated on the coated and uncoated wells for 1 h at room temperature. Using a multi-species conjugate (SBVMILK; obtained from ID Screen® Schmallenberg virus Milk Indirect ELISA; IDvet) diluted 1/80 for 1 h at room temperature detection was performed after the addition of tetramethylbenzidine (TMB) substrate (IDEXX) at a wavelength of 450 nm. After each step, the plates were washed three times with Tris-buffered saline with Tween 20. For readout, absorbances were calculated by subtracting the optical density (OD) measured on the uncoated wells from the values obtained from the protein-coated wells for each respective sample. Reproducibility was confirmed and normalization was achieved by reference to negative and positive sera samples.

#### Neutralization Assay

To evaluate specifically the presence of virus-neutralizing antibodies in serum samples we performed a virus neutralization test. Sera were pre-diluted (starting dilution from 1/16 to 1/512) with Dulbecco’s modified Eagle’s medium (DMEM) in a 96-well deep well master plate. 100 µL of this pre-dilution were transferred into a 96-well plate. A log2 dilution was conducted by passaging 50 µL of the serum dilution in 50 µL DMEM, leaving 50 µL of sera dilution in each well. Subsequently, 50 µL of SARS-CoV-2 (BavPat1) virus dilution (100 TCID_50_/well) was added to each well and incubated for 1 h at 37 °C. Lastly, 100 µL of trypsinized VeroE6 cells (cells of one confluent T-175 flask per 100 mL) in DMEM with 1% penicillin/streptomycin supplementation was added to each well. After 72 h incubation at 37 °C, the cells were evaluated by light microscopy for a specific CPE. A serum dilution was counted as neutralizing in the case no specific CPE was visible and is given as neutralizing dose 100 (ND100). The virus titer was confirmed by virus titration; positive and negative serum samples were included. Tests were performed in 3 technical replicates and average values were used to calculate the 100% neutralizing dose with the Kerber formula: (-log2) = a/b + c ((a) cell culture wells without virus replication, (b) number of cell culture wells per sera dilution, (c) -log2 of pre-dilution of the sera/yolk sample).

### Cell culture

#### Cell lines

Cells were obtained from Thermo Scientific if not stated otherwise. African green monkey kidney cells (Vero E6) were kindly provided by V. Thiel, Bern, Switzerland or obtained from the Collection of Cell Lines in Veterinary Medicine CCLV-RIE 0929. Adenocarcinomic human alveolar basal epithelial cells (A549) were obtained from NIBSC (A549-ACE-2 Clone 8-TMPRSS2; product number 101006). The THP-1 myelomonocytic leukemia cell line was obtained from the American Type Culture Collection. HEK293T cells were kindly provided by D. D. Pinschewer.

#### Maintenance

Cells were kept in DMEM high glucose with 10% FBS + 1% Penicillin/ Streptomycin for general propagation. DMEM high glucose was supplemented with 2% FBS + 1% Penicillin / Streptomycin for viral infection experiments. During the initial culture, the JAK-I inhibitor Pyridone 6 (CAS 457081-03-7) was added to a final concentration of 2µM as well as the NFκB inhibitor QNZ (CAS 545380-34-5) at 20nM. HEK293T-indE received in addition Doxycycline (Merck, D5207) to a final concentration of 2 µg/mL for induction.

#### Cell line generation

<u>HEK293T-E</u> were generated by transfecting HEK293T with 2µg plasmid DNA containing the SARS-CoV-2 E gene under CMV promoter control in a pcDNA3.1 background containing a Hygromycin resistance gene. After transfection cells were put in DMEM containing 250µg/mL of Hygromycin. The selection was kept for two weeks and clones were generated by limiting dilution before E expression was tested by RT-qPCR. The clone that showed the highest RNA expression levels was kept for downstream application.

<u>HEK293T-indE</u> (HEK293T-E Tet:E-IRES-ORF6) are a derivative of HEK293T-E with a second-generation lentiviral vector generated with the pCW57-E-IRES-ORF6 (Addgene plasmid #80921) as a transfer vector. The vector codes for SARS-COV-2 E and ORF6 under a Tetracycline inducible promoter. After infection, cells were selected in DMEM containing 20µg/mL of blasticidin for two weeks. Cells were analyzed by RT-qPCR for E and ORF6 induction following doxycycline treatment (fig. S1C).

<u>HEK293T-ACE2</u> were obtained by infecting the cells with a 2nd generation lentiviral vector with pHR-PGK_hACE2 (Addgene plasmid #161612) as a transfer vector. Cells were sorted for surface expression of ACE2 stained by Mouse anti-human ACE2 (R&D #FAB9332G).

<u>VeroE2T</u> were generated by transfecting Vero E6 cells with 2µg of an equimolar plasmid mixture containing the SARS-COV-2 E/ORF6/ORF7a/ORF8 genes in individual plasmids all under the CMV promoter in a pcDNA3.1 background containing a Hygromycin resistance gene. After transfection cells were cultivated in DMEM containing 250µg/mL of Hygromycin. Human TMPRSS2 expression in VeroE2T and in VeroE6 cells (<u>VeroE6-TMPRSS2</u>) was achieved by infecting the cells with a 2nd generation lentiviral vector pLEX307-TMPRSS2-blast (Addgene plasmid #158458) as a transfer vector. After infection cells were selected in DMEM containing 20µg/mL of blasticidin for two weeks and analyzed by RT-qPCR for transgene expression (fig. S2D).

### Plasmids and lentivirus

The genes of interest from the Wuhan strain (B.1) were inserted into the pcDNA3.1 backbone under the control of the CMV promoter for expression. The all-in-E plasmid contains the SARS-CoV-2 genes E and ORF6 under control of an ELF1α promoter or an IRES sequence, respectively, followed by ACE2 and TMPRSS2 under PGK promoter control separated by a P2A cleavage site in a pcDNA3.1 background. The integrity of all plasmids was verified by Sanger sequencing.

The plasmids required for the generation of second-generation lentiviruses were obtained from Addgene. Lentiviruses were generated by transfecting HEK-293T cells with pCMVR8.74 (RRID:Addgene_22036), pMD2G (RRID:Addgene_12259), and pLEX307-TMPRSS2-blast (RRID:Addgene_158458) plasmids. The culture medium was changed 5 hours after transfection, supernatant was collected 24 hours later and filtered through a 0.22µm filter to remove cellular debris.

### Genome reconstitution procedures for virus

Virus recovery was achieved as described in (22). In brief PCR fragments (fr A-D) spanning the whole SARS-CoV-2 genome were amplified using the high-fidelity proofreading enzyme Q5^®^ High-Fidelity DNA Polymerase (NEB, M0491L) in a 25 µL reaction volume using respective primers (fig. S1A, table S1).

Cycling conditions were used as recommended by the manufacturer. Fragments were obtained using the following primer combinations: frA: CMV for + frA-frB rev; frB: frB-frA for + frB-frC rev; frC: frC-frB for + frC-frD rev; frD: frD-frC for + SV40 rev. DNA oligonucleotides used are listed in table S1.

12-30 reactions were pooled and purified by PCR column purification using QIAquick PCR purification kit (Qiagen, 28104). DNA concentration was measured by Nanodrop 1000 (Thermo Fisher) or Quantus (Promega, QuantiFluor® ONE dsDNA System, E4871). DNA was further purified by ethanol precipitation and the final concentration was adjusted to 1 µg/µL in nuclease-free water.

Equimolar ratios of frA, frB, frC, frD or ΔfrD and all-in-E plasmid were transfected into HEK293T-indE using jetPRIME® (Polyplus, cat. 101000001) as recommended by the manufacturer. 4-24h post-transfection, medium was changed to DMEM 2% FBS with addition of JAK-I inhibitor Pyridone 6 (CAS 457081-03-7) to a final concentration of 2µM as well as the NFκB inhibitor QNZ (CAS 545380-34-5) at 20 nM and 2 µg/mL Doxycycline and Vero E2T were added for co-incubation. Every 3-4 days, the medium was exchanged. Screen for virus progeny production was done with SARS-CoV-2 antigen quick-test (Roche, 9901-NCOV-01G) (or CPE in E2T) and confirmed by RT-qPCR and FFA.

### Virus propagation for viral stocks

For *wild-type* controls, clinical isolates Muc-1 (a Wuhan-1-type virus isolate, provided by G. Kochs, University of Freiburg, Germany (SARS-CoV_Muc)), BavPat1 (SARS-CoV-2 Germany/BavPat1/2020, GISAID accession EPI_ISL_406862, kindly provided by Bundeswehr Institute of Microbiology, Munich, Germany), XBB.1.5 (isolated from nasopharyngeal aspirates of human donors, who had given their informed consent (approval by Ethikkommission Nordwest-und Zentralschweiz #2022-00303)), synthetic SARS-CoV-2 (Wuhan-1, GenBank No. MT108784 (*34*)) or rCoV2 (recombinant Wuhan-1-type virus produced by genome reconstitution (*22*), were propagated in VeroE6 cells until CPE was observed.

For deletion mutants, viral particles produced by HEK293T-indE were further amplified in VeroE2T cells, with additional trans-complementation of the all-in-E plasmid. Viral propagation was observed and monitored by CPE and Antigen quick-tests (*22*) and confirmed by RT-qPCR and FFA.

Final viral stocks were harvested, filtered by 0.2 µm filters to remove cells and frozen in small aliquots. For each viral stock, the viral titer was determined by RT-qPCR and FFA or titration by plaque forming assay.

All work including infectious SARS-CoV-2 viruses and its recombinant variants was conducted in a biosafety level 3 facility at the Department Biomedicine within the University of Basel (approved by the Swiss Federal Office of Public Health (BAG) #A202850/3).

### Standard plaque forming assay

*Wild-type* viral titers were determined by counting plaque-forming units (PFU) after incubation on susceptible cells. VeroE6 cells were seeded at a density of 4*10^6^ cells/96-well flat bottom plate in DMEM 2% FBS and incubated overnight at 37°C and 5% CO2. Virus was added 1:10 onto the cell monolayer in duplicates or triplicates and serially diluted 1:2 or 1:3. Plates were incubated for 2 days at 34°C, 5% CO_2_ until plaque formation was visible. For virus inactivation, 80μl of formaldehyde (15% w/v in PBS) (Merck, F8775) was added for 10 min to the cultures. After this period, fixative and culture medium were aspirated, and crystal violet (0.1% w/v) was added to each well and incubated for 5 min. Subsequently, the fixed and stained plates were gently rinsed several times with tap water and dried prior to analysis on a CTL ImmunoSpot® analyser.

### RNA extraction for viral quantification and sequencing of viral stocks

Viral RNA was extracted using the automated Promega Maxwell RSC system (Promega, AS4500) using either the Maxwell® RSC Viral Total Nucleic Acid Purification Kit (Promega, AS1330) or the Maxwell® RSC miRNA from the Tissue and Plasma or Serum Kit (Promega, AS1680).

### Sanger sequencing

The region of interest was amplified using SuperScript™ IV One-Step RT-PCR System (Thermo Fisher, 12594100) with either F-D2 IDRA4 or F-26847 and R-29046 N. The integrity of the PCR product was checked on agarose gel and subsequently sent for Sanger sequencing (for primers see table S1) to evaluate genome regions affected by deletions/mutations (Microsynth, Switzerland).

### Next-generation sequencing (NGS)

Viral RNA was converted to cDNA using cDNA Synthesis kit (biotechrabbit). cDNA was NGS sequenced using EasySeq SARS-CoV-2 WGS Library Prep Kit (NimaGen, SKU: RC-COV096) on an Illumina NextSeq 2000 system with a P1 flow cell (300 cycles). All NGS sequencing and raw data analysis was done by Seq-IT GmbH & Co. KG.

### RT-qPCR quantification of viral and intracellular RNA

For detection of SARS-CoV-2 RNA, a primer and TaqMan probe set for ORF-1b (table S1) were used as described (*37*). For the detection of SARS-CoV-2 E and TMPRSS2 an in-house primer / probe set was used (table S1). For normalization of mRNA expression GAPDH was used (table S1). For RT-qPCR Luna® Universal Probe One-Step RT-qPCR Kit (E3006E) was used according to manufactureŕs protocol. In brief Master Mix was set up: for one reaction 1 µL of each primer, 0.5 µL Probe, 10 µL of Luna Universal Probe One-Step Reaction Mix (2X), 1 µL of Luna WarmStart RT Enzyme Mix (20X) were mixed and brought to15 µL with nuclease free water. 15 µL of Master Mix were mixed with 5 µL RNA and amplified on ABI7500 fast cycler (ThermoFisher) using following cycling conditions: 10 min 55 °C, 1min 95°C denaturation, followed by 40 cycles for 10 seconds at 95°C and 30 seconds at 58°C.

### *In vitro* passaging for *in vitro* safety experiments

For viral passaging experiments, VeroE6 cells were infected with an MOI of 1 (based on FFU) for 3-4h with the *wild-type* or respective deletion candidate. The cells were then washed and fresh 2% DMEM medium was added. Every second day supernatant (SN) was passaged on freshly seeded VeroE6 (50% confluency). SNs for passage 1 (p1) and p2 were diluted 1:10, for all subsequent passages, SN was diluted 1:100. All collected passages p1 to p10 were subsequently passaged on VeroE2T. On day 3 and day 6 post infection SN was sampled for RT-qPCR and images of cell cultures were taken with a Leica DM IL LED inverted microscope. All conditions were treated equally.

### Biochemical procedures

For validation and comparison of vaccine candidate viruses, VeroE6+TMPRSS2 cells were infected with virus variants at an MOI of 0.1. 24h after infection, cells were washed twice with PBS before lysis in cold 140mM NaCl, 50mM Tris-HCL, 1% Triton-X100, 0,1%SDS, 0,1% sodium deoxycholate. supplemented with protease and phosphatase inhibitors (ThermoFisher, 1861281). Lysates were centrifuged for 10 min, 16’000g at 4°C and supernatants analyzed by Immunoblot. Signals were acquired using an image analyzer (Odyssey CLx, Licor).

### Flow cytometry analysis

#### Transfection

Cells were transfected using JetPrime (Polyplus, 101000001) transfection reagent according to the manufacturer’s protocol. Five hours after transfection, the culture medium was replaced. In the case of THP-1 cells, only ¼ of the recommended amount of DNA and reagents were used to avoid toxicity.

#### Infection

For cytometry experiments, all infections were conducted in DMEM supplemented with 2% FBS using a multiplicity of infection (MOI) value of 0.1 based on FFU (focus forming unit) data.

#### Staining

Cells were washed in PBS and stained with Zombie UV® Fixable Dead Cell Stain (Biolegend), and rinsed once with PBS and blocked in blocking buffer (PBS with 50% FCS, FcR Blocking Reagent 1:150 (Miltenyi Biotec) for 30 minutes at room temperature, followed by incubation with antibodies against cell-surface molecules in staining buffer (PBS with 15% FBS, FcR Blocking Reagent 1:1000) for 30 minutes at room temperature. Data were acquired on the Aurora (Cytek, Amsterdam, Netherlands) equipped with 5 lasers (355, 405, 488, 561, and 640 nm) and 60 channels (full spectrum cytometry), unmixed with SpectroFlo®, and analysed with FlowJo 10.0.7 (TreeStar).

### Immunocytochemistry

For detection of infectious vaccine viral particles (focus forming assay (FFA)), protein expression analysis and surface labeling, VeroE6-TMPRSS2 cells grown on coverslips in 24-well plates were infected with virus variants in 500 µL DMEM medium supplemented with 2% FCS and 1% Penicillin/Streptomycin and incubated overnight.

Cells were fixed with 4% PFA in PBS for 10 min at room temperature, washed and subsequently stained. For FFA and protein expression analysis, cells were blocked with 10% Normal Donkey Serum (Jackson ImmunoResearch, 017-000-121) and 0.1% Triton X-100 at room temperature for 60 min followed by incubation with primary antibodies for 60 min at room temperature or overnight at 4°C in 1% Normal Donkey Serum, 1% BSA and 0.3% Triton X-100 in PBS. Cells were washed three times for 10 min with 0.1% BSA / PBS and incubated with fluorophore-coupled secondary antibodies for 60 min at room temperature in 1% Normal Donkey Serum, 1% BSA and 0.3% Triton X-100 in PBS. Cells were washed once with 0.1% BSA / PBS and washed three times with PBS before mounting on microscope slides using Fluoromount-G (SouthernBiotech, 0100-01). For surface labeling, cells were blocked with 5% milk powder in PBS at room temperature for 1hr and incubated with primary antibodies in 1% BSA / PBS overnight at 4°C. After 3 washes with PBS, fluorophore-coupled secondary antibodies in 1% BSA / PBS were applied for 60 min at room temperature washed three times with PBS before mounting on microscope slides. Phalloidin-iFluor488 or -iFluor555 was co-applied with secondary antibodies to label F-actin (Abcam, ab176753 and ab176756 resp.). Hoechst 33342 dye (Merck, B2261) was co-applied during washing at a final concentration of 0.5 µg/mL for nuclear staining.

Images for FFA were acquired on a bright-field microscope (Nikon Ti2 equipped with a Photometrics 95B camera, Nikon NIS AR software), using a 20x Plan-Apochromat objective (numerical aperture 0.75) and were then processed in Fiji and Omero. For quantification of infected foci, images were analyzed with QuPath. Images for protein expression and surface labeling were acquired on an inverted spinning-disk confocal microscope (Nikon Ti2 equipped with a Photometrics Kinetix 25mm back-illuminated sCMOS, Nikon NIS AR software), using 40x and 100x Plan-Apochromat objectives (numerical aperture 0.95 and 1.45 respectively) and were then processed in Fiji and Omero.

### Antibodies

The following antibodies were used in this study: mouse monoclonal anti-β-actin (Cell Signaling Technology; 3700; RRID: AB_2242334; LOT# 20), rabbit polyclonal anti-SARS-CoV-2 nsp2 (GeneTex; GTX135717; RRID: AB_2909866; LOT# B318853), mouse monoclonal anti-SARS-CoV-2 Nucleocapsid protein (4F3C4, gift from Sven Reiche (*38*), sheep polyclonal anti-SARS-CoV-2 ORF3a (*39*), rat monoclonal anti-SARS-CoV-2 ORF6 (8B10, gift from Yoichi Miyamoto (*40*)), rabbit polyclonal anti-SARS-CoV-2 ORF8 (Novus Biologicals; NBP3-07972; LOT# 25966-2102), mouse monoclonal anti-SARS-CoV-2 Spike protein (4B5C1, gift from Sven Reiche).

Fluorophore-conjugated secondary antibodies were from Jackson ImmunoResearch (Cy3 donkey anti-rat #712-165-153, Cy3 donkey anti-mouse #715-165-151, Cy5 donkey anti-rabbit #711-175-152, Cy5 donkey anti-mouse #715-175-511), Li-Cor (IRDye 680RD donkey anti-mouse #926-68072, IRDye 680RD goat anti-rabbit #926-68071, IRDye 680RD goat anti-rat #926-68076) and Invitrogen (Alexa Fluor 647 donkey anti-mouse #A31571, Alexa Fluor 680 donkey anti-sheep #A21102).

Flow cytometry antibodies -all anti-human-were from Miltenyi REAfinity™ (VioBlue™ anti CD44 #130-113-344, VioGreen™ anti HLA-ABC #130-120-436, PerCP-Vio-700 anti CD59 #130-128-812, PE-Vio®770 anti CD275 (B7-H2) #130-116-805, APC anti CD70 # 130-130-100), Biolegend (Brilliant Violet 711 anti CD80 #305236, Alexa Fluor® 700 anti HLA-DR #307626) and R&D (mouse monoclonal anti-hACE2 #FAB9332G).

### Electron microscopy

Viral particles were fixed in 1% glutaraldehyde (Thermo Scientific, 233281000). A 4 µL aliquot of sample was adsorbed onto holey carbon-coated grid (Lacey, Tedpella, USA), blotted with Whatman 1 filter paper and vitrified into liquid ethane at -180°C using a Leica GP2 plunger (Leica microsystems, Austria). Frozen grids were transferred onto a Talos 200C Electron microscope (FEI, USA) using a Gatan 626 cryo-holder (GATAN, USA). Electron micrographs were recorded at an accelerating voltage of 200 kV using a low-dose system (40 e-/Å2) and keeping the sample at – 175°C. Defocus values were -2 to 3 µm. Micrographs were recorded on 4K x 4K Ceta CMOS camera.

### Quantification and statistical analysis

Statistical analysis was conducted with GraphPad Prism 9. Sample sizes were chosen based on previous experiments and literature surveys. No statistical methods were used to pre-determine sample sizes. Appropriate statistical tests were chosen based on sample size and are indicated in individual experiments.

## Acknowledgments

We are grateful to Laurent Perez and Christian Münz for helpful comments on the manuscript. The authors thank the BioEM lab and Mohamed Chami of the Biozentrum, University of Basel, for their support with electron microscopy and the DBM Microscopy Core Facility for support with image acquisition and analysis. We thank Sven Reiche (FLI, Greifswald, GER) for the generous provision of monoclonal anti-S and anti-N antibodies, which were obtained through the EU H2020 project EVA-GLOBAL, project #871029, and Yoichi Miyamoto (Natl. Inst. Biomed. Innovation, Health and Nutrition, Osaka, J) for the anti-ORF6 antibody. We are thankful to Mareen Grawe, Angele Breithaut and Tobias Britzke and the animal caretakers for their essential help. We thank G. Kochs, Freiburg, GER for providing the SARS-CoV-2 Wuhan isolate and Martin Daeumer, Alex Thielen (Seq-It GmbH, Kaiserslautern, GER) and Adrian Egli & team (USB, Basel, CH) for expert NGS support. pCMVR8.74 and pMD2.G were a gift from Didier Trono (RRID:Addgene_12259), pLEX307-TMPRSS2-blast was a gift from Alejandro Chavez & Sho Iketani (RRID:Addgene_158458). pHR_PGK-hACE2 was a gift from Brad Rosenberg (Addgene plasmid # 161612 ; http://n2t.net/addgene:161612 ; RRID:Addgene_161612). Vero E6 cells and synthetic SARS-CoV-2 virus (Wuhan-1, GenBank No. MT108784) were kindly provided by V. Thiel. Figures were created with BioRender.com

## Author contributions

This work was jointly conceived by M.J.L., F.O., D.H., M.B., C.M. and T.K., experimental procedures were performed by M.J.L, F.O., D.H., J.S., E.K., N.J.H., L.U., Y.Z., C.W., L.U., D.Ho. and C.L., data analysis was conducted by M.J.L., F.O., D.H. and J.S.. The manuscript was jointly written by M.J.L, F.O., D.H. and T.K., with editing provided by J.S., D.Ho., M.B. and C.M.. All authors have read and approved the final version of the manuscript.

## Data and materials availability

Further information and requests for resources and reagents should be directed to and will be fulfilled by the lead contact, Thomas Klimkait. Plasmids will be deposited at Addgene.

