## Supplementary Material for "Single-cycle SARS-CoV-2 vaccine elicits high protection and sterilizing immunity in hamsters"

**The supplementary PDF file includes:**

Figs. S1 to S5
Tables S1 to S5

**Other Supplementary Materials for this manuscript include the following:**

Movie S1

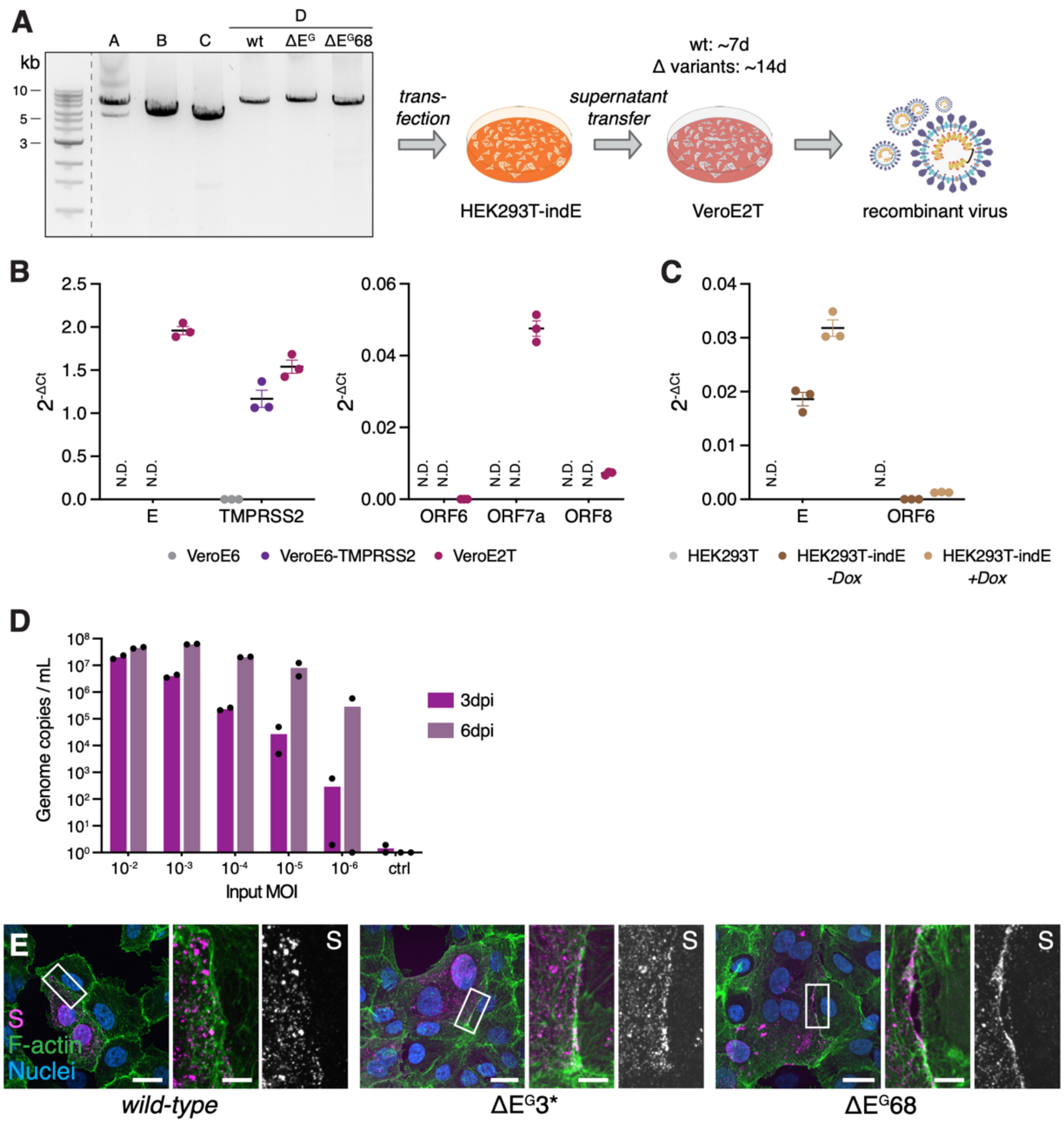


**Figure S1**

(A) Scheme showing virus production. Representative DNA gel showing PCR products of the four fragments A, B, C and D (for D: wild-type (wt), $\Delta E^G$ and $\Delta E^G68$) amplified from plasmids encoding individual fragments A-D of SARS-CoV-2 followed by transfection of HEK293T-indE cells for spontaneous genome reconstitution and supernatant transfer onto VeroE2T cells for virus harvest.

(B) Quantification of transgenes by RT-qPCR for expression of E and TMPRSS2 (left panel) or ORF6, ORF7a and ORF8 (right panel) relative to GAPDH in VeroE6, VeroE6-TMPRSS2 or VeroE2T cells, non-detectable signal annotated with N.D. (n=3 independent mRNA isolations).

(C) Quantification of transgenes by RT-qPCR for expression of E and ORF6 relative to GAPDH in HEK293T or HEK293T-indE treated +/- doxycycline for 48h, non-detectable signal annotated with N.D. (n=3 independent mRNA isolations).

(D) Infection of complementing VeroE2T cells with $\Delta E^G$ at different MOIs and analysis by RT-qPCR of the ORF1b NSP14 regions normalized to external SARS-CoV-2 standards after 3- and 6-days post infection (n=2 infected cultures).

(E) Surface labeling of S (magenta), F-actin (green), and nuclei (blue) in VeroE6-TMPRSS2 cells infected with wild-type SARS-CoV-2, $\Delta E^G$, or $\Delta E^G68$.

Mean and S.E.M, scale bar in (E) is 20$\mu$m for overview and 5$\mu$m for ROI images.

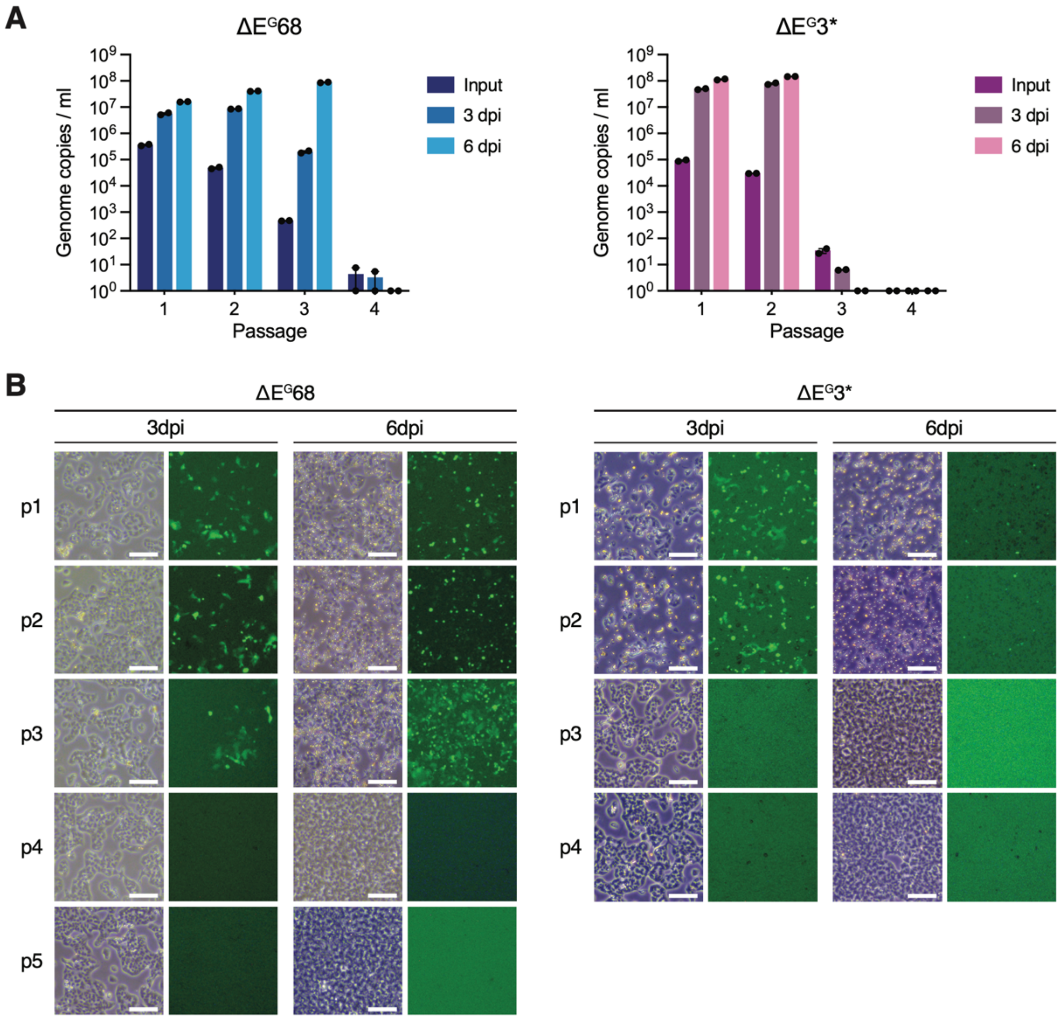


**Figure S2.**
(A) Analysis of supernatants from VeroE6 passaging experiment on complementing VeroE2T cells by RT-qPCR, passages 1-4 from ΔE$^{G}$68 and ΔE$^{G}$3* were analyzed at timepoints 0 (Input) or 3- and 6-days post infection (n=2 technical replicates).
(B) Images of infected, complementing VeroE2T cells with supernatants from passaging experiment (p1-p4 or p5) after 3- and 6-days post infection for ΔE$^{G}$68 and ΔE$^{G}$3*. Images show bright field view on the left, expression of eGFP on the right.
Mean and S.E.M, scale bar in (B) is 200$\mu$m.

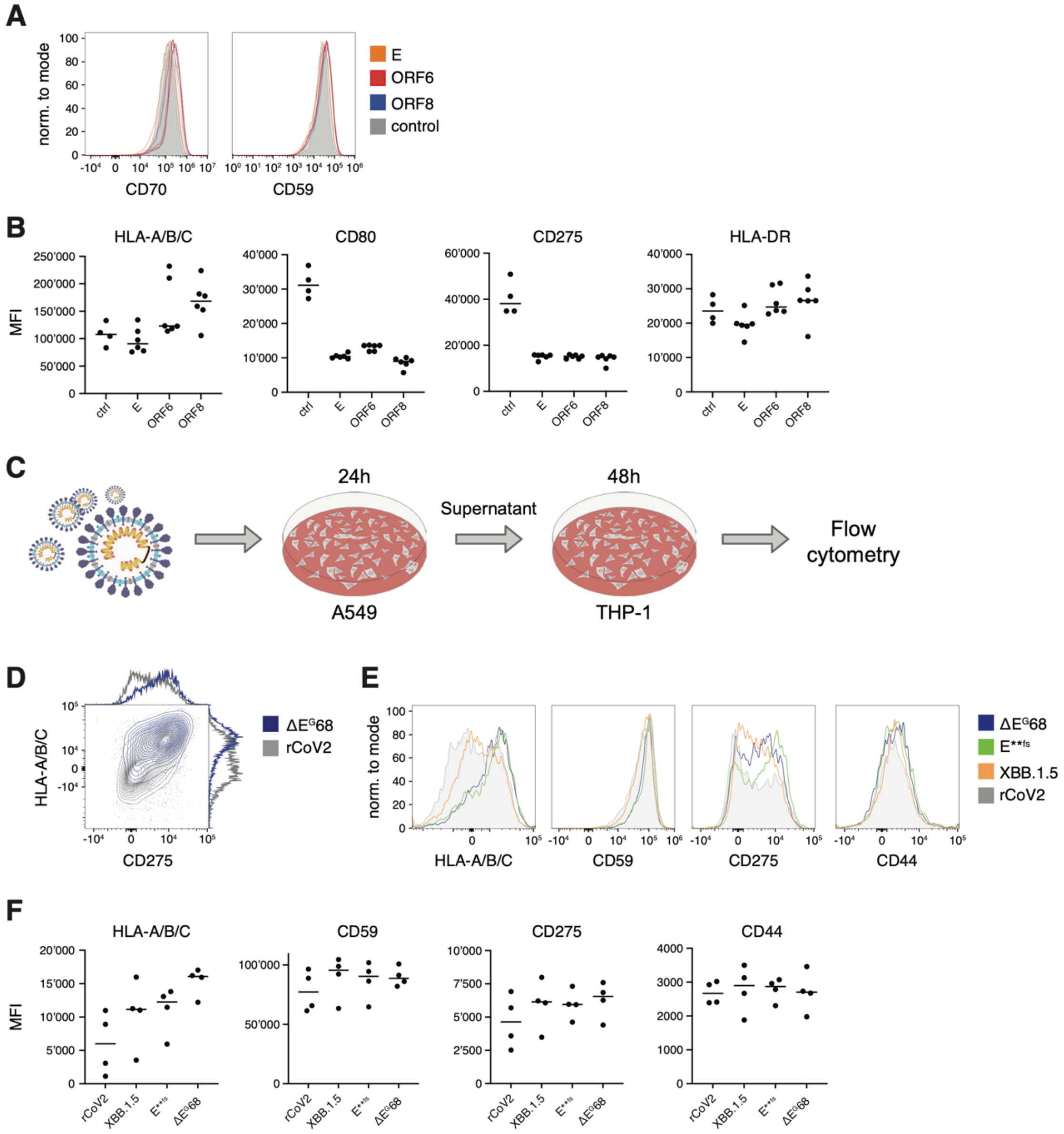


**Figure S3.**
(A-B) Modulation after transfection: (A) Flow cytometry staining for surface expression of CD70 and CD59 on THP-1, 48h after transfection with plasmids coding for ORF6, ORF8, or Envelope proteins, compared with control transfection. (B) Median fluorescence intensity of HLA-A/B/C, CD275, CD80, and HLA-DR corresponding to the main Fig 3.
(C-F) Modulation after infection: (C) Method representation: HEK-293T-ACE2-TMPRSS2 cells were infected with rCoV2, E**fs, $\Delta E^{G}68$, or XBB.1.5 SARS-CoV-2 virus (MOI = 0.1) for 24h and their respective supernatant was applied on THP-1 for 48h before surface staining and analysis (Fig. 3). (D) Contour plot comparing the expression of HLA-A/B/C and CD275 on the HEK-293T. (E) Histogram showing the expression of HLA-A/B/C CD59, CD275, and CD44 on HEK-293T. (F) Median fluorescence intensity of HLA-A/B/C, CD59, CD275, and CD274 for the different replicates of the experiment.

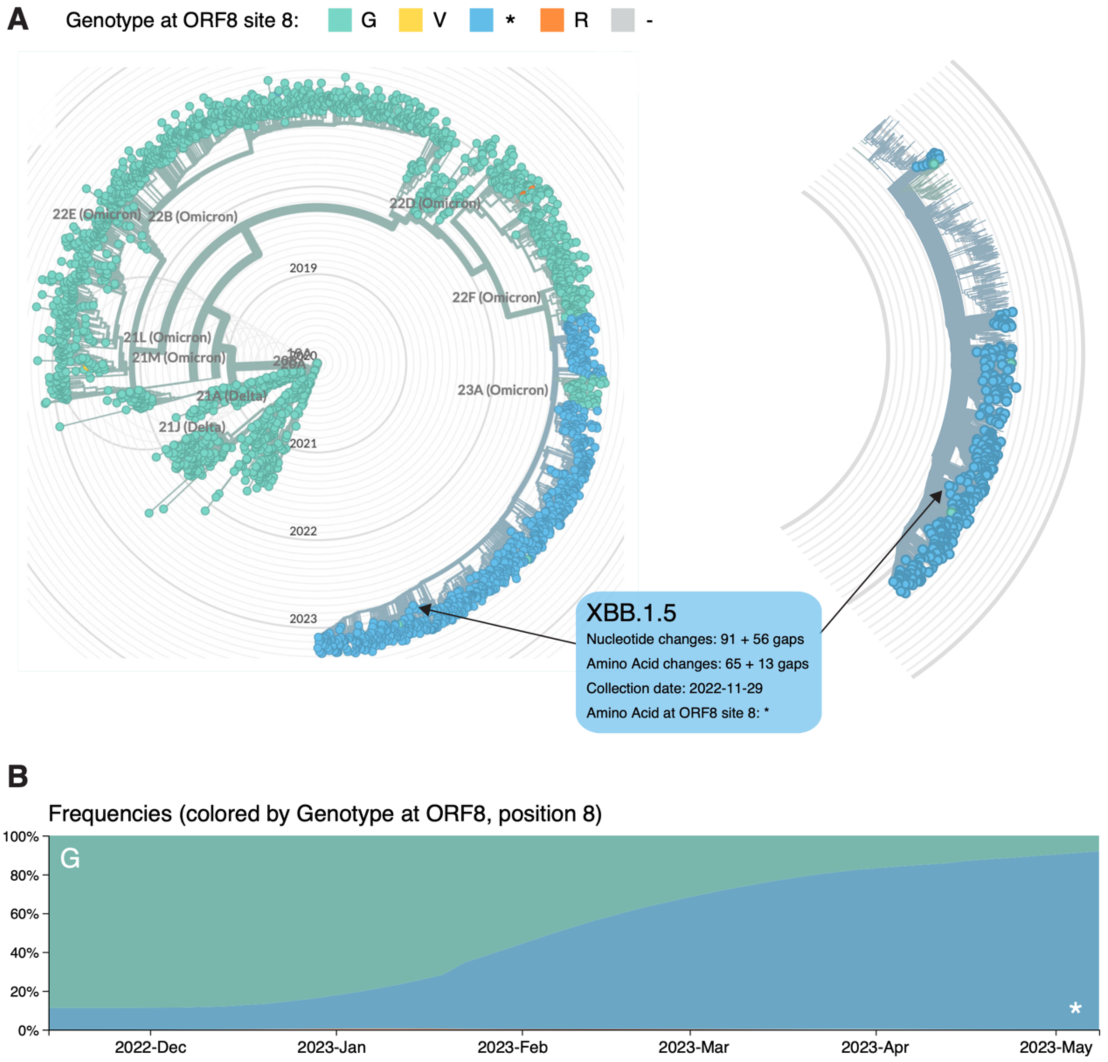


**Figure S4.**

(A) Circular phylogenetic tree of SARS-CoV-2 virus isolates from the currently ongoing SARS-CoV-2 pandemic. Left: Shown are 2711 representative genomes from GISAID color coded for amino acid 8 of ORF8 and Clade name next to each tree branch. Isolines show time of isolation from initial occurrence of SARS-CoV-2. Amino acid color code is shown in legend next to plot. Data update from 2023-05-06. Source: https://nextstrain.org/ncov/gisaid/global/6m

Right: Snippet of circular phylogenetic tree highlighting all XBB.1.5 SARS-CoV-2 variants of concern (n=371) carrying mainly a premature stop codon at position 8 of ORF8. All features of XBB.1.5 are annotated for a representative isolate (indicated with arrow).

(B) Frequency of SARS-CoV-2 isolates from (A) having a Glycine or a stop codon at position 8 of ORF8 over the period of 6 months (Nov 2022 – May 2023).

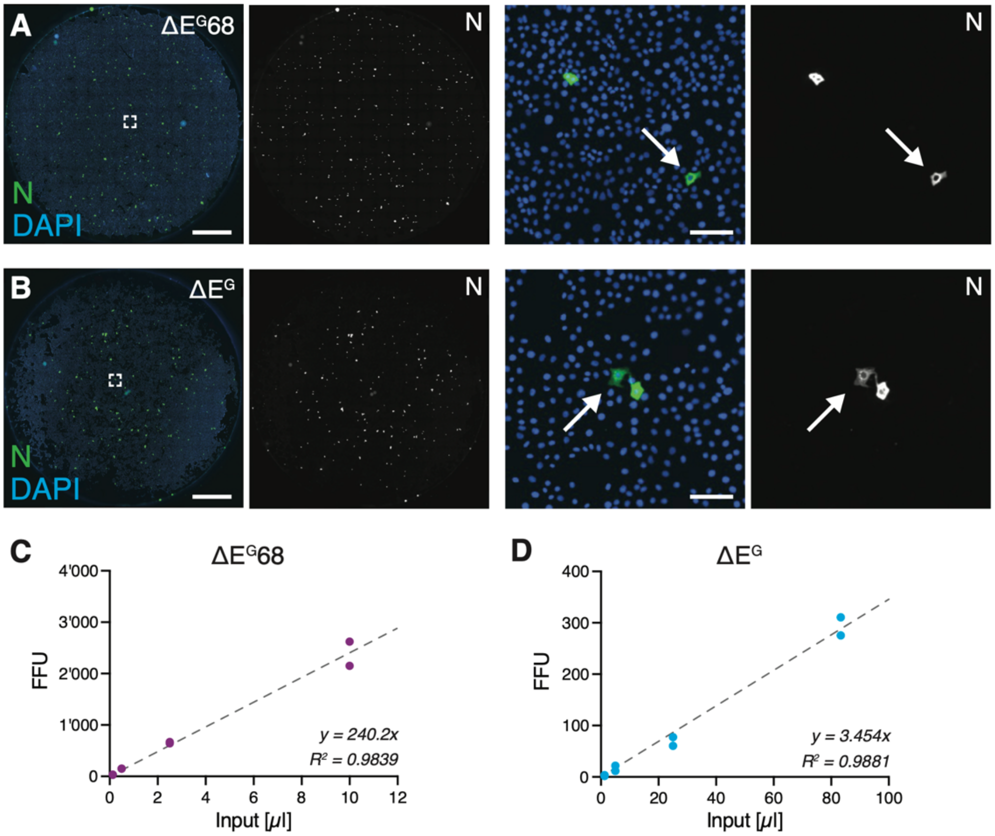


**Figure S5.**
(A, B) Representative example of infected cells used to quantify viral titers of SCVs on coverslips in a 24-well culture dish for $\Delta E^G68$ (A) and $\Delta E^G$ (B) in VeroE6-TMPRSS2 cells, detection of N shown in green, nuclei are stained with Hoechst (blue). Left: overview images, right: region of interest images showing individual infected cells as indicated.
(C, D) Titration of $\Delta E^G68$ (C) or $\Delta E^G$ (D) and quantification by FFA (n=2). Linear fit and correlation indicated for determination of SCV titers used to inoculate Syrian hamsters.
Scale bar is 2mm and 100$\mu$m for both (A) and (B) (overview and ROI images resp.).

**Table S1.**

Methods Supplement: Oligonucleotide primers and qPCR probes.

| Name (gene / fragment) | Sequence |
|---|---|
| E-Sarbeco for | ACA GGT ACG TTA ATA GTT AAT AGC GT |
| E-Sarbeco rev | ATA TTG CAG CAG TAC GCA CAC A |
| E-Sarbeco probe | FAM-ACA CTA GCC ATC CTT ACT GCG CTT CG-BHQ-1 |
| nCoV IP4-14059 for | GGT AAC TGG TAT GAT TTC G |
| nCoV IP4-14146 rev | CTG GTC AAG GTT AAT ATA GG |
| nCoV IP4-14084 probe | FAM-TCA TAC AAA CCA CGC CAG G-BHQ-1 |
| β-actin for | CAG CAC AAT GAA GAT CAA GAT CAT C |
| β-actin rev | CGG ACT CAT CGT ACT CCT GCT T |
| β-actin probe | HEX-TCG CTG TCC ACC TTC CAG CAG ATG T-BHQ1 |
| ORF1b for | TGG GGT TTT ACA GGT AAC CT |
| ORF1b rev | AAC ACG CTT AAC AAA GCA CTC |
| ORF1b probe | FAM-TAG TTG TGA TGC AAT CAT GAC TAG-BHQ1 |
| Envelope for | GCG TAC TTC TTT TTC TTG CTT TCG |
| Envelope rev | TTG CAG CAG TAC GCA CAC AA |
| Envelope probe | FAM-CAC TAG CCA TCC TTA CTG CGC TTC GA-BHQ1 |
| ORF6 for | GCA GAG ATA TTA CTA ATT ATT ATG AGG ACT TTT A |
| ORF6 rev | TCT CCA TTG GTT GCT CTT CA |
| ORF6 probe | FAM-TCC ATT TGG AAT CTT GAT TAC ATC ATA AAC CTC A-BHQ1 |
| ORF7a for | CGA GGG CAA TTC ACC ATT TC |
| ORF7a rev | CGT GTT TTA CGC CGT CAG GA |
| ORF7a probe | FAM-TGC ACT GAC TTG CTT TAG CAC TCA ATT TGC-BHQ1 |
| ORF8 for | CCT TTA ATT GAA TTG TGC GTG GA |
| ORF8 rev | CCC AAT TTA GGT TCC TGG CAA |
| ORF8 probe | FAM-TGA GGC TGG TTC TAA ATC ACC CAT TCA GT-BHQ1 |
| TMPRSS2 for | CTC TAA CTG GTG TGA TGG CG |
| TMPRSS2 rev | TGC CAG GAC TTC CTC TGA G |
| TMPRSS2 probe | FAM-CGG ACC AAA CTT CAT CCT TCA GG-BHQ1 |
| GAPDH for | GAA GGT GAA GGT CGG AGT C |
| GAPDH rev | GAA GAT GGT GAT GGG ATT TC |
| GAPDH probe | FAM-CAA GCT TCC CGT TCT CAG CC-BHQ1 |
| CMV for | CGA TGT ACG GGC CAG ATA TAC G |
| frA-frB rev | GTG TTA TTA AAT AGA AAA TAG CAG CAA CAA AAA GGA ACA CAA GTG TAA CTT TAA TTA ACT GCT TCA ACC |

| Name (gene / fragment) | Sequence |
|---|---|
| frB-frA for | GCA CTT AAG GGT GGT AAA ATT GTT AAT AAT TGG TTG AAG CAG TTA ATT AAA GTT ACA CTT GTG TTC C |
| frB-frC rev | AAA CTG TCT ATT GGT CAT AGT ACT ACA GAT AGA GAC ACC AGC TAC GGT GCG AGC TCT ATT CTT TGC AC |
| frC-frB for | TAT AAC TCA AAT GAA TCT TAA GTA TGC CAT TAG TGC AAA GAA TAG AGC TCG CAC CGT AGC TGG TG |
| frC-frD rev | ATC ACC AAT CAA AGT TGA ATC TGC ATC AGA GAC AAA GTC ATT AAG ATC TGA GTC GAC AAG CAG CG |
| frD-frC for | TAC AGC TGT TTT AAG ACA GTG GTT GCC TAC GGG TAC GCT GCT TGT CGA CTC AGA TCT TAA TGA CTT TGT C |
| SV40 rev | GCG GCC GCC AGA CAT GAT AAG |
| D2 for | GGA ACT GTA ACT TTG AAG CAA GGT G |
| 29046 N rev | CGA CGT TGT TTT GAT CGC GCC C |
| 26847 for | GGA ACC AAT TTA TGA TGA ACC GAC G |
| 26526 SARS2 La for | GCA GAT TCC AAC GGT ACT ATT ACC |
| M 574 for | TGT GAC ATC AAG GAC CTG CC |

**Table S2.**

Envelope qPCR analysis of nasal washings of $\Delta E^{G}68$ vaccinated animals to test for *wild-type* reversion in vaccinated animals or contact animals at different time points. The contact animal that tested positive for SARS-CoV-2 antibodies is highlighted in grey. Values are Ct values measured with SARS-CoV-2 Envelope or beta-actin specific primers. For negative controls, plain PBS was used and processed along with nasal washings (N.D.: not detected).

| | ID | dpim 0 | | dpim 3 | | dpim 7 | | dpim 24 | |
|---|---|---|---|---|---|---|---|---|---|
| | | E | β-actin | E | β-actin | E | β-actin | E | β-actin |
| $\Delta E^{G}68$ | 1 | N.D. | 35.71 | N.D. | 31.87 | N.D. | 32.17 | N.D. | 30.58 |
| $\Delta E^{G}68$ | 2 | N.D. | 34.25 | N.D. | 31 | N.D. | 30.57 | N.D. | 32.45 |
| $\Delta E^{G}68$ | 3 | N.D. | N.D. | N.D. | 31.35 | N.D. | 30.19 | N.D. | 29.25 |
| $\Delta E^{G}68$ | 4 | N.D. | 34.66 | N.D. | 29.87 | N.D. | 32.17 | N.D. | 30.52 |
| $\Delta E^{G}68$ | 5 | N.D. | 31.64 | N.D. | 31.92 | N.D. | 30.13 | N.D. | 30.71 |
| $\Delta E^{G}68$ | 6 | N.D. | 33.29 | N.D. | 33.36 | N.D. | 32.65 | N.D. | 32.82 |
| $\Delta E^{G}68$ | 7 | N.D. | 35.94 | N.D. | 29.08 | N.D. | 32.09 | N.D. | 29.66 |
| contact | 8 | N.D. | 35.5 | N.D. | 33.83 | N.D. | 31.79 | N.D. | 30.32 |
| $\Delta E^{G}68$ | 9 | N.D. | N.D. | N.D. | 33.16 | N.D. | 32.74 | N.D. | 33.33 |
| contact | 10 | N.D. | 34.55 | N.D. | 31.23 | N.D. | 28.44 | N.D. | 31.07 |
| $\Delta E^{G}68$ | 11 | N.D. | 37.15 | N.D. | 31.65 | N.D. | 29.18 | N.D. | 30.31 |
| contact | 12 | N.D. | 35.64 | N.D. | 33.44 | 39.27 | 33.84 | N.D. | 28.53 |
| $\Delta E^{G}68$ | 13 | N.D. | 34.66 | N.D. | 30.73 | N.D. | 31.06 | N.D. | 31.93 |
| contact | 14 | N.D. | 35.24 | N.D. | 32.79 | N.D. | 30.53 | N.D. | 28.79 |
| $\Delta E^{G}68$ | 15 | N.D. | 38.35 | N.D. | 32.24 | N.D. | 31.45 | N.D. | 30.68 |
| contact | 16 | N.D. | 37.58 | N.D. | 32.38 | N.D. | 29.06 | N.D. | 29.04 |
| $\Delta E^{G}68$ | 17 | N.D. | 39.58 | N.D. | 28.61 | N.D. | 30.15 | 39.2 | 30.79 |
| contact | 18 | N.D. | N.D. | N.D. | 32.2 | N.D. | 28.76 | N.D. | 30.62 |
| negative control | 1 | N.D. | N.D. | N.D. | N.D. | N.D. | N.D. | N.D. | 36.48 |
| | 2 | N.D. | 38.62 | N.D. | N.D. | N.D. | 36.93 | N.D. | 39.43 |
| Muc-1 (B.1) $10^{-1}$ | 1 | 19.79 | 35.16 | | | | | | |
| | 2 | 19.69 | 34.51 | | | | | | |
| Muc-1 (B.1) $10^{-2}$ | 1 | 22.87 | 36.36 | | | | | | |
| | 2 | 22.95 | N.D. | | | | | | |
| Muc-1 (B.1) $10^{-3}$ | 1 | 26.13 | 38.07 | | | | | | |
| | 2 | 26.22 | N.D. | | | | | | |
| Muc-1 (B.1) $10^{-4}$ | 1 | 30.03 | N.D. | | | | | | |
| | 2 | 29.96 | N.D. | | | | | | |

**Table S3.**

Serum neutralization data for $\Delta E^{G}68$ vaccinated animals and contacts after vaccination and challenge infection against Wuhan (B.1). Note: all values represented with “<dilution” are negative for the indicated dilution or higher dilutions but are not tested for lower dilutions. Values represent mean from 3 technical replicates, calculated with Kerber formula (see Methods).

| | ID | dpim 19 | dpim 33 | dpc 5 | dpc 14 |
|---|---|---|---|---|---|
| $\Delta E^{G}68$ | 1 | <1:32 | 1:203.2 | 1:128 | |
| $\Delta E^{G}68$ | 2 | <1:32 | 1:322.5 | 1:161.3 | |
| $\Delta E^{G}68$ | 3 | <1:64 | 1:406.4 | 1:128 | |
| $\Delta E^{G}68$ | 4 | <1:32 | <1:32 | 1:512 | |
| $\Delta E^{G}68$ | 5 | <1:32 | 1:645.1 | 1:322.5 | |
| $\Delta E^{G}68$ | 6 | <1:32 | 1:101.6 | 1:128 | |
| $\Delta E^{G}68$ | 7 | <1:32 | 1:80.6 | | 1:161.3 |
| contact | 8 | | | | |
| $\Delta E^{G}68$ | 9 | <1:32 | 1:161.3 | | 1:64 |
| contact | 10 | | | | |
| $\Delta E^{G}68$ | 11 | <1:32 | 1:256 | | 1:101.6 |
| contact | 12 | | | | |
| $\Delta E^{G}68$ | 13 | <1:32 | 1:161.3 | | 1:128 |
| contact | 14 | <1:32 | 1:40.32 | | <1:32 |
| $\Delta E^{G}68$ | 15 | <1:32 | <1:32 | | 1:256 |
| contact | 16 | | | | |
| $\Delta E^{G}68$ | 17 | <1:32 | 1:406.4 | | 1:128 |
| contact | 18 | | | | |

**Table S4.**

Serum neutralization data for ΔE$^{G}$ vaccinated animals, contacts and sham-treated animals after vaccination and challenge infection against Wuhan (B.1). Note: all values represented with “<dilution” are negative for the indicated dilution or higher dilutions but are not tested for lower dilutions. Values represent mean from 3 technical replicates, calculated with Kerber formula (see Methods).

| | ID | dpim 19 | dpim 33 | dpc 5 | dpc 14 |
|---|---|---|---|---|---|
| ΔE$^{G}$ | 1 | <1:128 | <1:128 | 1:512 | |
| ΔE$^{G}$ | 2 | <1:128 | N.A. | 1:256 | |
| ΔE$^{G}$ | 3 | <1:128 | <1:128 | | 1:322,5 |
| contact | 4 | | | | 1:40,32 |
| ΔE$^{G}$ | 5 | <1:512 | <1:256 | 1:101,6 | |
| ΔE$^{G}$ | 6 | <1:128 | <1:32 | 1:128 | |
| ΔE$^{G}$ | 7 | <1:512 | <1:256 | | 1:50,8 |
| contact | 8 | | | | 1:32 |
| ΔE$^{G}$ | 9 | <1:256 | <1:256 | 1:1024 | |
| ΔE$^{G}$ | 10 | <1:512 | <1:256 | | 1:512 |
| contact | 11 | | | | 1:40,32 |
| sham | 12 | | | <1:16 | |
| sham | 13 | | | 1:20,16 | |
| sham | 14 | | | <1:16 | |
| sham | 15 | | | <1:16 | |

**Table S5.**

IP4 qPCR analysis of nasal washings of $\Delta E^{G}68$ vaccinated and contact animals after miscalculated challenge infection on days 1, 2, and 3 after challenge, measured with IP4 and β-actin primer / probe sets. Inoculum: Ct=35.64, 1089 gc/ml (N.D.: not detected).

| | ID | dpc 1 | | dpc 2 | | dpc 3 | |
|---|---|---|---|---|---|---|---|
| | | IP4 | β-actin | IP4 | β-actin | IP4 | β-actin |
| $\Delta E^{G}68$ | 1 | N.D. | 33.84 | N.D. | 31.60 | N.D. | 26.96 |
| $\Delta E^{G}68$ | 2 | N.D. | 30.65 | N.D. | 30.70 | N.D. | 28.79 |
| $\Delta E^{G}68$ | 3 | N.D. | 32.11 | N.D. | 32.34 | N.D. | 29.53 |
| $\Delta E^{G}68$ | 4 | N.D. | 30.75 | N.D. | 32.38 | N.D. | 30.94 |
| $\Delta E^{G}68$ | 5 | N.D. | 29.68 | N.D. | 31.82 | N.D. | 31.09 |
| $\Delta E^{G}68$ | 6 | N.D. | 30.96 | N.D. | 30.93 | N.D. | 30.35 |
| $\Delta E^{G}68$ | 7 | N.D. | 29.46 | N.D. | 29.07 | N.D. | 31.58 |
| contact | 8 | N.D. | 29.77 | N.D. | 32.02 | N.D. | 29.79 |
| $\Delta E^{G}68$ | 9 | N.D. | 31.77 | N.D. | 31.37 | N.D. | 31.11 |
| contact | 10 | N.D. | 32.99 | N.D. | 32.42 | N.D. | 29.15 |
| $\Delta E^{G}68$ | 11 | 38.06 | 30.87 | N.D. | 31.26 | N.D. | 27.90 |
| contact | 12 | N.D. | 34.12 | N.D. | 32.69 | N.D. | 31.35 |
| $\Delta E^{G}68$ | 13 | N.D. | 31.38 | N.D. | 33.76 | N.D. | 29.01 |
| contact | 14 | N.D. | 28.32 | N.D. | 30.01 | N.D. | 27.91 |
| $\Delta E^{G}68$ | 15 | N.D. | 31.74 | N.D. | 29.47 | N.D. | 29.59 |
| contact | 16 | N.D. | 32.17 | N.D. | 31.15 | N.D. | 28.61 |
| $\Delta E^{G}68$ | 17 | N.D. | 30.97 | N.D. | 31.31 | N.D. | 32.10 |
| contact | 18 | N.D. | 31.95 | N.D. | 30.44 | N.D. | 29.61 |

**Movie S1.**

Summary of the single-cycle vaccine concept
